# Functional genomics guided multi-omics framework identifies aldehyde metabolism as a therapeutic vulnerability in Fanconi anaemia

**DOI:** 10.64898/2026.08.26.747186

**Authors:** Khalid Saeed, Bader Ahmari, Hemza Ghadbane, Caroline Heckman, Ziaurrehman Tanoli

## Abstract

Understanding the molecular vulnerabilities associated with Fanconi anemia (FA) is essential for identifying therapeutic opportunities and elucidating the mechanisms underlying disease progression and cancer predisposition. However, progress in this area remains constrained by limited availability of representative FA cellular models. To address this challenge, we defined an “FA-like” cellular state by identifying cancer cell lines exhibiting high-dependency on core FA pathway genes, and integrated CRISPR-Cas9 gene essentiality data at multiple molecular layers, including mutation, copy number alterations, mRNA expression, and independent patient-derived transcriptomic datasets. Functional enrichment analyses highlighted biological pathways previously implicated in FA pathogenesis, most notably aldehyde detoxification, cholesterol/fatty acid metabolism, and androgen signaling. Analysis of LINCS-L1000 perturbational transcriptomics resource identified compounds, capable of reversing the FA-associated transcriptional signature, further supporting the pharmacological tractability of the identified molecular vulnerabilities. In addition, drug-target affinity analysis prioritized aldehyde-metabolizing enzymes, including ALDH1A1 and ALDH2, as potentially druggable candidates. Notably, disulfiram demonstrated predicted high-affinity interactions with multiple proteins involved in aldehyde and lipid metabolism, including ALDH1A1, ALDH2, and MGLL, supporting its potential for further investigation in FA-related settings. Although additional validations are required, the identified vulnerabilities and candidate targets provide a foundation for future mechanistic and therapeutic investigations in FA and FA-associated malignancies.

## INTRODUCTION

Fanconi anemia (FA) is a rare genetic disorder characterized by bone marrow failure (BMF), developmental abnormalities, and increased susceptibility to cancer. The condition arises from defects in the repair of DNA interstrand crosslinks (ICLs)^1^. The FA pathway consists of at least 23 core FA proteins and several FA-associated proteins (FAAPs), functioning in a sequential, multistep process. Loss-of-function mutations in any of the FA genes can lead to constitutional genomic instability and disrupt additional cellular pathways that may contribute to malignancies^2,3^. Allogeneic hematopoietic stem cell transplantation (HSCT) remains the standard treatment for FA-associated BMF and can restore hematopoiesis. However, HSCT does not correct the underlying genetic defect in non-hematopoietic tissues and therefore is not fully curative ^4^.

Despite the fundamental DNA repair defect, spontaneous somatic mosaicism occurs in a subset of patients through genetic reversion or other compensatory events in hematopoietic stem and progenitor cells (HSPCs), giving rise to blood cell populations with partially restored FA pathway function ^5^. These naturally occurring rescue mechanisms demonstrate that FA-associated defects can be functionally compensated. In support of this concept, recent genetic perturbation and large-scale loss-of-function screening studies have identified suppressor interactions and compensatory pathways capable of alleviating DNA repair deficiencies beyond the canonical FA pathway. For example, disruption of the BLM helicase complex was shown to suppress FA complementation group (FANC) C (FANCC)-associated phenotypes, an interaction that was subsequently validated in FA complementation group D2 (FANCD2)-deficient cells ^6^. Similarly, inactivation of the deubiquitylating enzyme USP48 enhanced DNA repair capacity and reduced chromosomal instability in FA-defective cells ^7^. Beyond direct modulation of DNA repair pathways, metabolic compensation has also emerged as a critical protective mechanism in FA. Enhanced aldehyde detoxification was recently shown to markedly reduce genotoxic stress and restore normal differentiation of hematopoietic progenitor cells lacking a functional FA pathway ^8^. Consistent with this observation, ALDH2-mediated detoxification of endogenous aldehydes has been demonstrated to protect against BMF, chromosomal instability, and hematopoietic stem cell attrition in FA models ^9^. In parallel, CRISPR-Cas9-based genome editing has demonstrated the potential to restore FA-associated cellular defects through correction of pathogenic mutations or the generation of compensatory genetic alterations ^10^. Despite these advances, the development of safe and effective gene-editing therapies remains challenging, highlighting the need to identify alternative therapeutic vulnerabilities and molecular targets ^11^.

Although cancer cell lines do not fully recapitulate the physiological context of FA, they may provide scalable systems for investigating genetic dependencies associated with FA pathway dysfunction. Such models help overcome the limitations of patient-derived FA samples, which are scarce, heterogeneous, and often clinically fragile ^12–14^.

Furthermore, large-scale CRISPR-Cas9 screening efforts have enabled systematic identification of context-specific vulnerabilities across diverse cellular backgrounds, offering new opportunities to uncover pathways that modulate sensitivity to FA related DNA repair defects (https://depmap.org/portal/ ^15,16^. However, gene dependency data alone provide limited insight into the molecular mechanisms underlying selective vulnerabilities. Integrating dependency profiles with complementary omics features, including gene expression, somatic mutations, and copy number alterations (CNA), may reveal biomarkers and pathways associated with FA-related cellular states. Conversely, perturbational transcriptomic resources such as the LINCS-L1000 dataset provide an additional opportunity to identify compounds that can reverse disease or phenotype-associated transcriptional signatures ^17^, thereby linking molecular vulnerabilities to potentially actionable pharmacological interventions.

In this study, we integrated FA gene dependency profiles from DepMap with multiple publicly available omics resources to define an FA-like cellular state and characterize its associated molecular dependencies. By combining CRISPR-Cas9 essentiality data with transcriptomic and genomic features, we sought to identify candidate vulnerabilities, uncover pathways associated with FA pathway dysfunction, and generate hypotheses for future mechanistic and therapeutic investigations. We further capitalized the LINCS-L1000 perturbational transcriptomics resource to identify compounds predicted to reverse the FA-associated transcriptional signature, providing an additional pharmacological layer for prioritizing potentially actionable vulnerabilities. Complementing this transcriptional rescue analysis, drug-target affinity analysis prioritized several aldehyde-metabolizing enzymes, including ALDH1A1 and ALDH2, as potentially druggable candidates. Although previous studies have used CRISPR dependency profiling, transcriptomic analyses, and perturbational signatures to investigate cancer vulnerabilities and drug responses, an integrated framework combining genome-wide dependency profiles with multi-omics features, patient-derived transcriptomic data, and compound-induced transcriptional responses has not, to our knowledge, been systematically applied to identify compensatory therapeutic vulnerabilities associated with FA and FA-associated malignancies.

### MATERIAL AND METHODS

### Cell line, CRISPR dependencies and multi-omics data

The data was compiled from multiple complementary lines of evidence, including genomic and transcriptomics profiles of cancer cell lines, obtained from publically available resources ^15,16^. The pooled CRISPR-Cas9 knockout screening data available through DepMap portal (DepMap Public 24Q4, https://depmap.org/portal/), which provides Chronos gene dependency scores across a panel of 1,178 cancer cell lines. In parallel, molecular datasets were collected for the corresponding cell lines, including mRNA expression (n = 1,597), copy number variation (CNV; n = 1,929), and damaging mutation profiles (n = 1,929). To ensure direct integration across data modalities, only cell lines with matched CRISPR dependency data and corresponding molecular profiles were retained for downstream analyses, resulting in a final dataset of 1,108 cell lines. In addition, DepMap portal provides multi-dose drug response data for approximately 5,000 compounds, which were leveraged for target identification. To determine whether the predicted drug-target interactions translated into phenotypic responses, we evaluated disulfiram sensitivity using the PRISM Repurposing dataset from DepMap, comprising 468 cancer cell lines, to assess whether the predicted molecular interactions were reflected in experimentally measured disulfiram sensitivity across diverse cancer cell lines.

### Data integration and preprocessing

All datasets were harmonized by matching common cell lines (DepMap; ACH-identifier) to ensure consistency across data modalities. Based on FA pathway-gene dependency scores quantified using Chronos (gene effect values), cell lines were categorized into three dependency classes: Low, Medium, and High, as illustrated in **Figure 2**. Cell lines with low Chronos scores, corresponding to high (FA pathway) dependency, were referred throughout the manuscript as ‘FA-like’ cell lines, as they exhibited increased sensitivity and reduced growth following CRISPR-Cas9-mediated knockout of FA genes.

**Figure 1:**
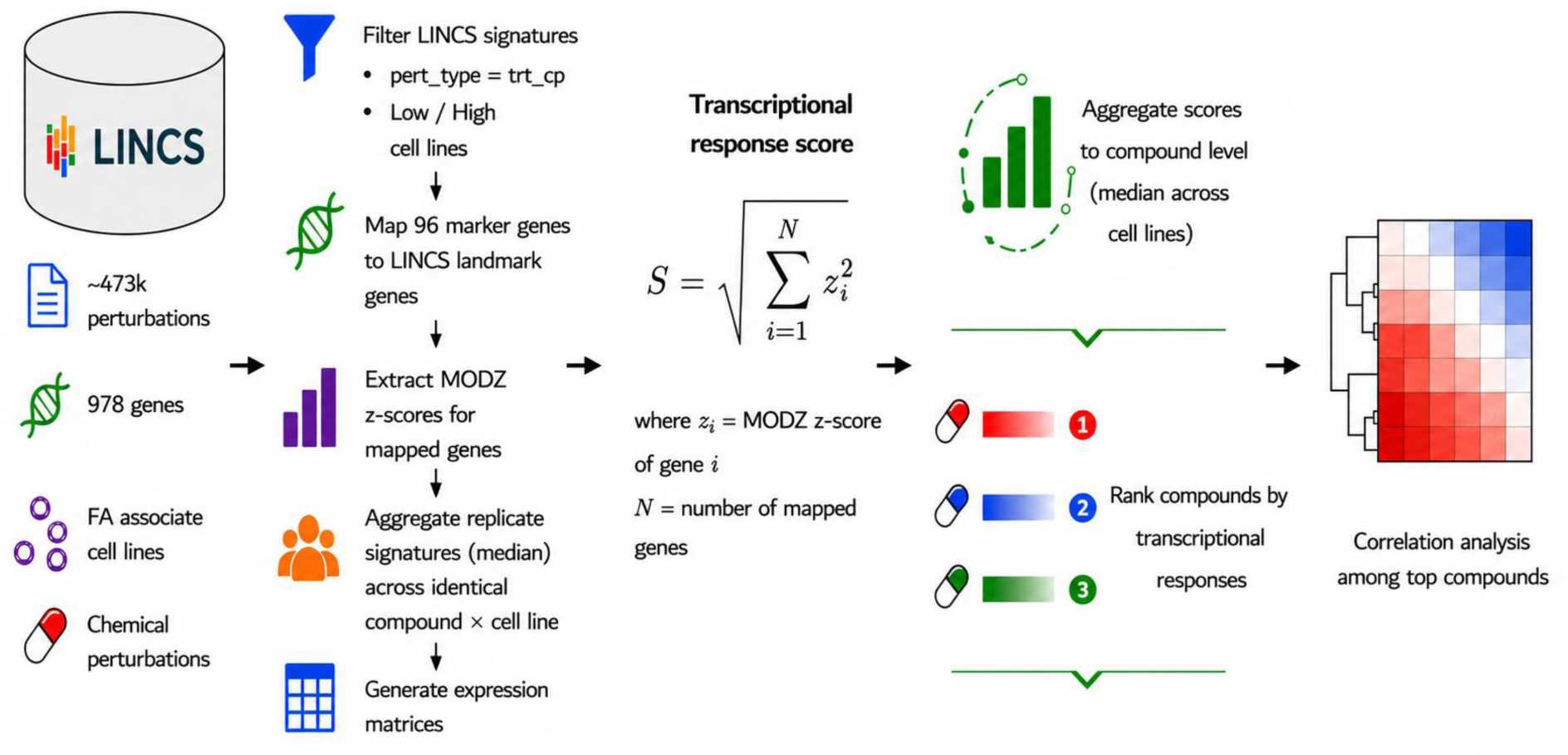
Workflow for identifying transcriptionally responsive compounds using LINCS L1000 perturbation profiles. LINCS L1000 Level 5 (MODZ) chemical perturbation data were used as the input for transcriptomic analysis. The preprocessing pipeline first filtered LINCS signatures to retain only chemical perturbations (pert_type = trt_cp) from the predefined FA-associated cell lines. The selected 96 FA-associated marker genes were then mapped to the LINCS landmark gene set, MODZ z-scores were extracted for the mapped genes, replicate perturbation signatures corresponding to identical compound-cell line pairs were aggregated using the median, and four expression matrices (collapsed, low-dependency, high-dependency, and Low − High) were generated. For each compound signature, a transcriptional response score was calculated as the Euclidean norm of the standardized marker gene expression profile. Compound-level scores were subsequently aggregated across cell lines using the median, compounds were ranked according to their transcriptional response scores, and the highest-ranking compounds were selected for downstream analyses. Finally, pairwise correlation analysis was performed among the top responsive compounds to identify compounds exhibiting similar transcriptional response patterns.

**Figure 2:**
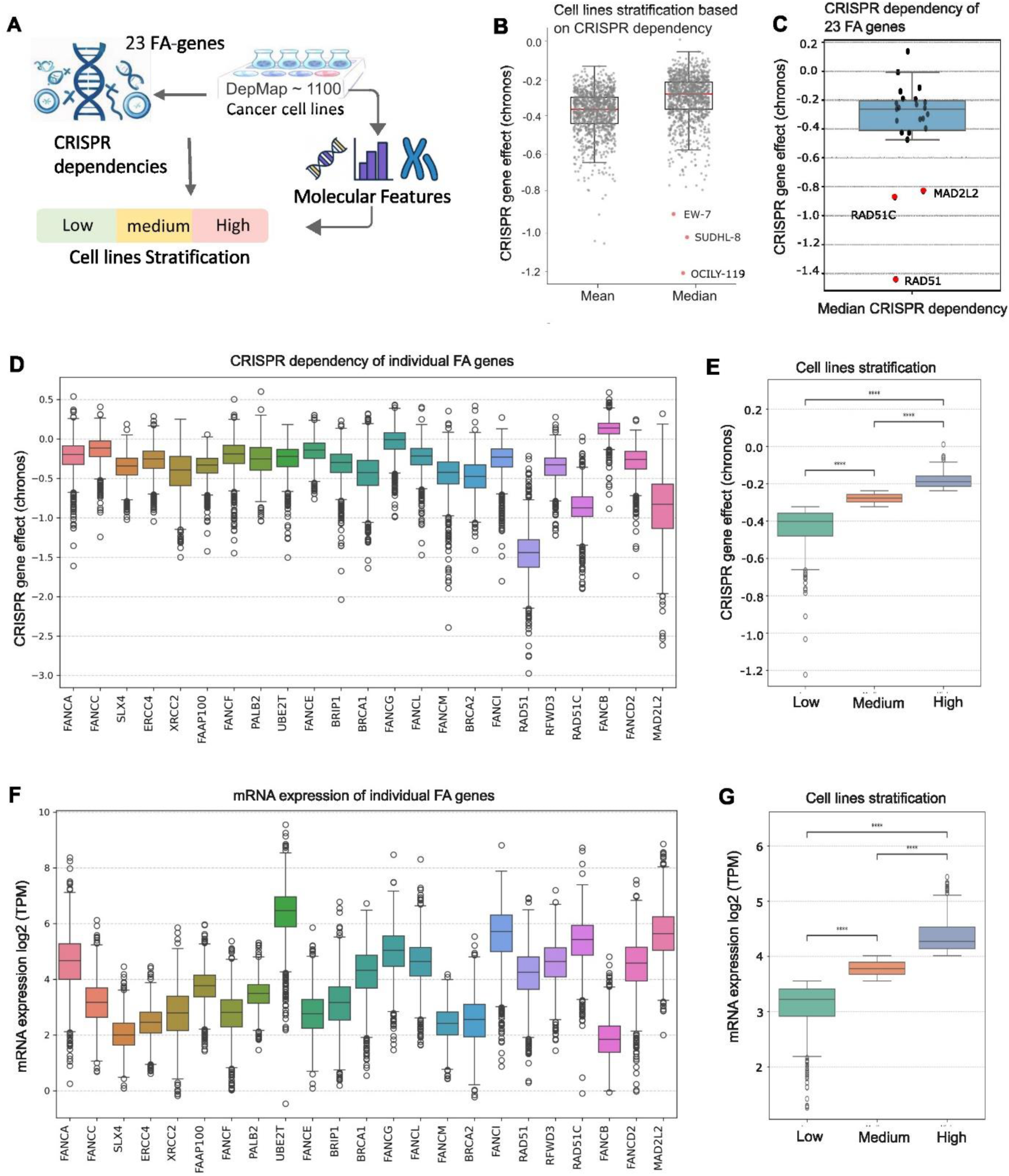
Identification of FA-mimicking cell models. **A)** Schematics of work flow to identify cancer cell stratification based on CRISPR dependencies and molecular profiles. **B)** The boxplot illustrates the combined dependency/features of 23 FA genes across a panel of 1178 cancer cell lines from diverse tissue origins, analysed using the DepMap portal (https://depmap.org/portal), each dot represent a cell lines, red dot represents the most vulnerable cell lines with lower CRISPR dependency group hence may referred to most FA-like cell models. **C)** plotted 23 FA genes, each dot represents individual FA genes, red dot represents the most essential genes whose absence is lethal to most of the cell lines. D) Individual FA genes CRISPR score across all cell lines **E)** combine, with stratification of cell lines as low, medium, and high dependency scores. CRISPR dependency, a value of −1.0 indicates strong gene dependency, comparable to known essential genes, whereas a score of 0 indicates no observable effect from gene knockout. **F)** Individual FA genes mRNA expression levels across all cell lines. **G)** Combined, with stratification of cell lines as low, medium, and high mRNA expressed cell lines. Statistical significance was determined using t-test. Significance levels are denoted as P < 0.05 (*), P < 0.01 (**), and P < 0.001 (***); ns = not significant.

### Multi-omics comparative and correlation analyses of FA gene dependency

To characterize molecular differences among dependency groups, mRNA expression, CNV, and mutation profiles were systematically compared across cell lines. The molecular stratification was visualized using annotated plots (**Figure 2, Supplementary Figure 1**), with features showing statistically significant differences highlighted to facilitate biological interpretation. This analysis enabled the identification of candidate genes and molecular features, potentially associated with FA gene dependency.

Associations between FA pathway gene dependency and multi-omics features, including mRNA expression, somatic mutations, and copy number variations (CNVs), were evaluated separately for each dependency group using pearson correlation analyses (**Supplementary Figure 2)**.

### Predictive modeling and feature ranking

Predictive modelling was performed using Random Forest (RF), XGBoost (XGB), and SHapley Additive explanations (SHAP) to identify molecular features, predictive of FA pathway CRISPR dependencies. More details on model development are shown in the section S1 of **Supplementary material**. The objective of the predictive modeling framework was to prioritize human genes according to their contribution to the FA-like phenotype, as inferred from cancer cell line molecular profiles. Cancer cell lines were categorized into three dependency groups (Low, Medium, and High) based on their FA pathway gene dependency (Chronos) scores. Multi-omics profiles, including mRNA expression, copy number variation (CNV), damaging mutations, and genetic interaction-derived features, were used as predictor variables, whereas the dependency category served as the response variable.

### Differential multi-omics analysis of FA dependency

To comprehensively characterize molecular differences associated with FA dependency, all statistically tested features (mRNA expression, mutations, and CNVs) were subsequently retained irrespective of the magnitude of their fold change as shown in the **Supplementary Figure 4**. Following Student’s *t*-test analysis, median, mean, standard deviation, fold change (High versus Low dependency groups), log2 fold change (log2FC), and *P*-values were calculated for each feature. Features with median values below 0.1 in both groups were excluded to minimize the influence of low-abundance variables. A volcano plot (**Supplementary Figure 4A**) was generated to visualize differential features by plotting log2 fold change against the negative logarithm of the *P*-value (−log10 *P*). This approach enabled visualization of the complete distribution of molecular alterations associated with FA dependency.

Next, we refined the analysis by applying more stringent criteria to the low- and high-dependency groups, thereby maximizing biological contrast and facilitating the identification of candidate biomarkers and compensatory mechanisms underlying FA pathway vulnerability. Thus, only samples representing the extreme phenotypes were retained for downstream analysis, with dependency scores < −0.5 classified as the high FA dependency group) and scores > −0.2 classified as the low FA dependency group). Similarly, fold change was calculated as the ratio of the median value in the high group relative to the low group, and log2 fold change (log2FC) was subsequently computed. Features with median values below 1.0 in both groups were excluded to minimize the influence of low-abundance variables (**Supplementary Table 2**). Only features exhibiting a median fold change >1.5 or <0.67 were retained as shown in the **Figure 4A**.

### Gene expression analysis of Fanconi anemia patient samples

To investigate transcriptional alterations associated with FA, we reanalyzed publicly available gene expression microarray data generated from bone marrow cells of healthy volunteers and FA-patients ^18^. The data were obtained from the Gene Expression Omnibus (GEO) under accession GSE16334. Differential gene expression analysis was performed by comparing FA samples with healthy controls to identify genes exhibiting altered expression in the disease state.

Differentially expressed genes were ranked according to their log fold change (logFC) and statistical significance. Consistent with the original analysis, FA and control bone marrow samples exhibited widespread transcriptional differences. For downstream comparative analyses, genes with an absolute log fold change (logFC) ≥ 1 were retained, corresponding to at least a two-fold difference in expression between FA and control samples. Statistical significance was assessed using the limma framework with false discovery rate (FDR) correction for multiple testing, and only genes meeting the predefined significance criteria were included for subsequent analyses (**Supplementary Figure 4B**).

Bulk RNA-seq data from Zubicaray et al., comprised mesenchymal stromal cells (MSCs) isolated from seven untreated patients with FA, all carrying pathogenic Fanconi anemia complementation group A (*FANCA)* variants, together with three healthy donor-derived MSC samples (BioProject: PRJNA1114677) ^19^. We also reanalyzed a single-cell RNA-seq dataset (GSE157591) of hematopoietic stem and progenitor cells (HSPCs) from seven patients with FA and five healthy donors reported by ^20^. Detailed analytical procedures are described in the **Supplementary Material section: S2.**

To assess the robustness of our findings, we compared concordant genes identified from the cancer cell line analyses with three independent patient-derived datasets. Despite differences in sample type and sequencing platform, all datasets demonstrated consistent dysregulation of multiple prioritized genes. For prioritization, we primarily relied on the bone marrow microarray dataset reported by Vanderwerf *et al.* because its tissue source most closely reflects the BMF phenotype central to our study. In contrast, the remaining datasets were generated from cultured MSCs ^19^ or enriched HSPCs ^20^, which may introduce cell type-specific transcriptional biases. Nevertheless, these independent datasets provided important orthogonal validation by supporting the directional concordance of our prioritized gene signatures.

### Identifying biomarker genes associated with Fanconi anemia

To identify robust candidate genes associated with FA dependency, we integrated feature selection results from multiple complementary analytical approaches. The top 200 ranked (∼1% genes) molecular features from each of the predicting algorithms (Random Forest classifier, gradient-boosted decision tree (XGBoost), and SHAP from **Figure 3**) analyses were combined with 684 differentially expressed mRNA features identified by univariate *t*-test analysis of RNA expression using predefined log fold change and statistical significance thresholds (**Figure 4A**). As the downstream analyses focused on transcriptional alterations, only RNA-derived features were retained. Duplicate genes identified by multiple methods were removed, resulting in a non-redundant set of (n = 1006) unique RNA features (**Supplementary table 3**).

**Figure 3:**
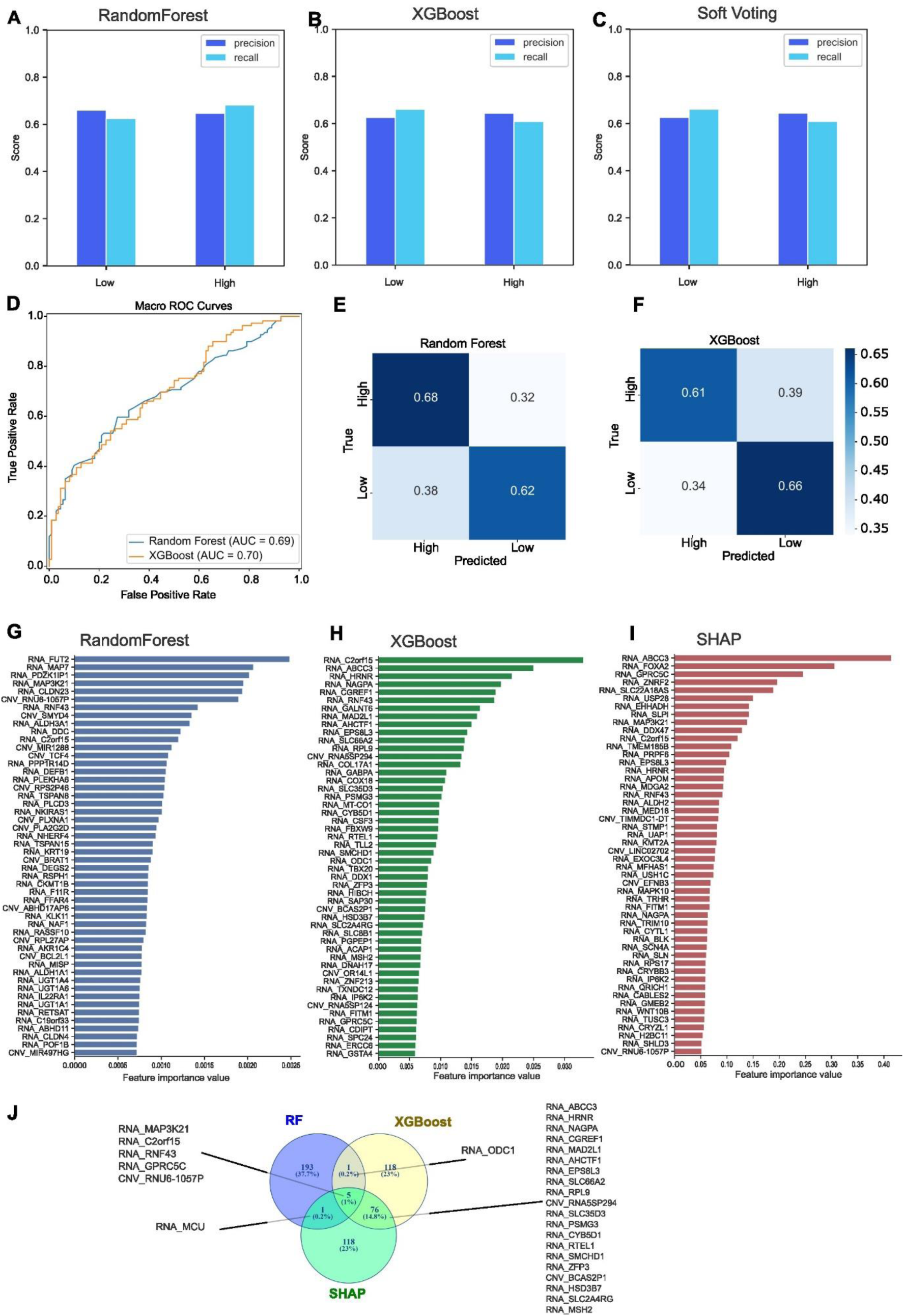
Identification of key determinants of CRISPR-based gene dependencies in low and high dependency group using supervised machine learning approach. Precision and recall for each dependency class obtained from these models using the independent testing dataset for **A)** Random Forest (RF), **B)** XGboost (XGB) classifier and **C)** soft voting ensemble. **D)** Receiver Operating Characteristic (ROC) curve was computed and plots the true positive rate (sensitivity) against the false positive rate (specificity) based on predicted class probabilities (Low, high dependency groups), and performance was summarized using Area Under the Curve (AUC), with higher values indicating improved classification. **E)** Confusion matrices evaluated per-class accuracy summarizes the number of true positives, false positives, false negatives, and true negatives for each class by comparing predicted labels against true labels for RF, **F)** XGB. **G-I)** Molecular features were ranked according to their contribution to model prediction using the intrinsic feature importance scores calculated by each algorithm. The top-ranking variables are shown in descending order of importance. **G)** Random Forest importance reflects the average reduction in Gini impurity across all decision trees. **H)** XGBoost importance represents the cumulative contribution of each feature during gradient-boosted tree construction. **I)** Mean absolute SHAP values were calculated from the trained XGBoost model using the independent testing dataset. Larger SHAP values indicate variables exerting greater influence on model predictions irrespective of direction. Features are ranked from highest to lowest overall contribution. **J)** Venn-diagram shows overlapping genes from the top 200 scoring features identified in each predicting model.

**Figure 4:**
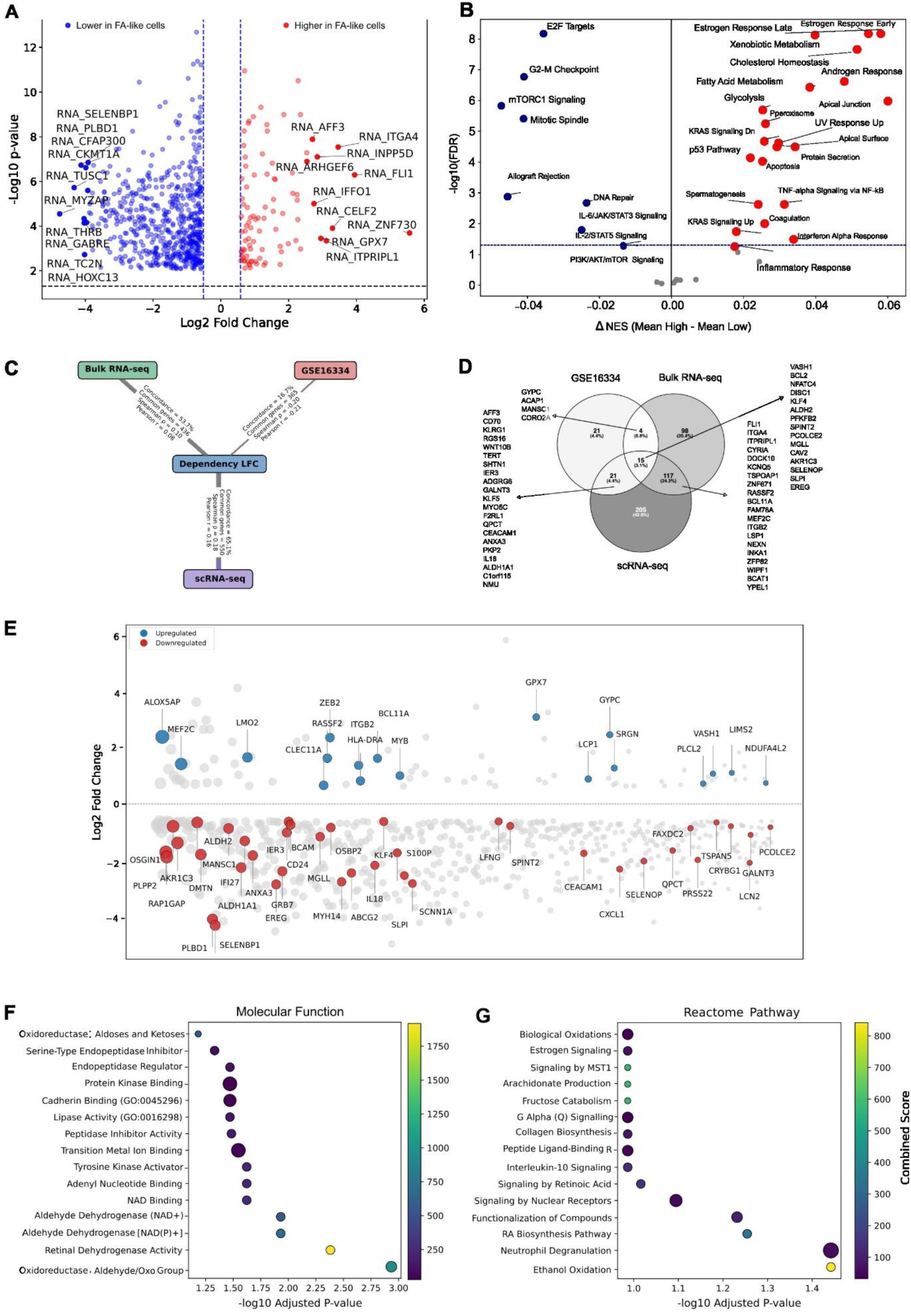
Transcriptomic signatures associated with Fanconi anemia-like CRISPR dependency. **A)** Volcano plot represents top ranked 684 differential mRNA expression between low vs high CRISPR dependent groups. The plot using the data only from the cell lines those dependency scores < −0.5 (high FA dependency) and > −0.2 (low FA dependency). Features with median values below 1.0 in both groups were excluded, and only feature with median fold change >1.5 or <0.67 were retained. **B)** Pathway enrichment score (per-sample pathway activities) using GSEApy based on differential expression of gene across low and high CRISPR dependencies (obtain from the top 200 features were selected from each of the top rank features of the RF, XGBoost, and SHAP analyses, while 684 features were identified by t-test based on RNA expression constitute a total of 1006 unique mRNA). **C)** Nodes represent the DepMap CRISPR dataset, GEO microarray, bulk RNA-seq, and single-cell RNA-seq datasets. Edges connect pairwise comparisons and are annotated with the percentage of concordant genes, the number of shared genes, Pearson’s correlation coefficient (r), and Spearman’s rank correlation coefficient (ρ). Edge thickness is proportional to the concordance percentage. **D)** Venn-diagram shows overlapping concordant genes of 3 independent datasets, and lists of common genes. **E)** Scatter plot showed differentially expressed 684 mRNA between low vs high CRISPR dependent groups. Blue and red highlighted genes are indicating those differentially expressed mRNA from FA patients vs healthy Bone marrow samples (Limma LFC cutoff +/-0.5) dot size indicating corresponding significance measured with FDR values. **F-G)** 42 concordant lower expressed genes were submitted to web-based gene-set enrichment analysis platform Enrichr (Ma’ayan Lab) against **F)** molecular function, **G)** Reactome.

To prioritize genes with potential clinical relevance, this combined feature set was further compared with differentially expressed genes identified from primary bone marrow samples of patients with FA (GEO accession: GSE16334) as shown in used for **Figure 4C**. This was performed by examining genes exhibiting differential expression between the lower and upper FA dependency quantiles in the cell line analysis and determining whether they were also differentially expressed in patient samples. This comparison identified 365 overlapping genes, of which 61 (highlighted in **Figure 4C**) displayed concordant directionality of expression i.e., consistently upregulated or downregulated in both the cell line models and patient-derived samples relative to their respective controls.

Because differential expression estimates originated from independent datasets generated using different experimental platforms and normalization procedures, comparisons were based on the direction of gene regulation rather than the absolute magnitude of fold change. Concordance of expression direction was evaluated using a binomial sign test and further assessed by direction-weighted spearman correlation (**Supplementary table 4**). Finally, the highest-ranked differentially expressed genes demonstrating concordant regulation across both datasets were selected for downstream gene-drug interaction analyses to identify candidate small molecules with predicted binding affinity toward the corresponding protein products.

### Identifying drug candidates associated with FA associated gene targets

To characterize pharmacological interactions involving the curated set of overlapping gene targets, we queried the ChEMBL database (v-36, latest available version at the time of analysis). ChEMBL is a comprehensive manually curated repository of experimentally validated bioactive molecules and drug-target interactions ^21^. ChEMBL was selected because it provides standardized bioactivity measurements collected from the published literature and other public sources, enabling systematic identification of approved drugs with known binding affinities toward human protein targets. Gene symbols corresponding to the nominated genes were programmatically mapped to their respective human UniProt identifiers to ensure compatibility with ChEMBL’s target annotation framework and minimize ambiguity arising from gene aliases or alternative protein names.

At the time of analysis, ChEMBL(v-36) contained experimentally validated binding affinity data of 4,005 approved drugs (including different approved formulations and salts) interacting with thousands of human protein targets. Since many approved drugs exhibit substantial polypharmacology and interact with numerous molecular targets, comprehensive retrieval of all drug-target relationships required large-scale database queries. Preliminary attempts to retrieve these data through the ChEMBL web API proved computationally inefficient and unstable for the required query volume. Therefore, the complete ChEMBL v36 PostgreSQL database dump (https://ftp.ebi.ac.uk/pub/databases/chembl/ChEMBLdb/latest/) was downloaded and deployed locally, providing a reproducible, scalable, and efficient framework for large-scale drug-target extraction and downstream analyses.

Drug-target interactions were filtered using stringent inclusion criteria to retain only high-confidence pharmacological associations. Specifically, only interactions satisfying all of the following criteria were included: (i) approved drugs (max_phase=4 in ChEMBL); (ii) single human protein targets; (iii) experimentally derived binding assay measurements; (iv) standardized potency values reported as pChEMBL, representing the negative logarithm of molar activity values (e.g., IC₅ ₀, Ki, Kd, EC₅ ₀, or AC₅ ₀); and (v) potent interactions defined by pChEMBL ≥ 5, corresponding to an activity threshold of ≤10 μM. Restricting the analysis to experimentally validated binding assays and standardized pChEMBL values ensured consistent comparison of drug potency across different studies and assay types while minimizing variability introduced by heterogeneous experimental conditions.

After applying the above filtering criteria, the resulting dataset comprised 2,031 approved drugs with experimentally validated interactions involving 1,689 human protein targets. This curated drug-target interaction network served as the reference pharmacological space for subsequent drug prioritization. The predefined resistant and sensitive-associated gene subsets were then intersected with the curated list of approved drug targets to identify existing therapeutics capable of modulating the nominated genes. This analysis identified 12 approved drugs (**Figure 6A**) with high-confidence experimentally validated binding to at least one of the 61 genes of interest, representing potential candidates for drug repurposing and further biological or clinical investigation.

### Drug response measurements in cell lines

To evaluate whether target engagement translated into phenotypic drug response, we queried NCI-60 for publicly available drug response measurements corresponding to the 12 identified approved drugs in **Figure 6**. Drug sensitivity data were examined across 63 lymphoid cell lines and 786 non-lymphoid cell lines (**Supplementary File 2**). Drug responses were assessed using pGI50 values, where higher values indicate increased growth inhibition (drug sensitivity), and lower values indicate relative resistance. In addition, drug sensitivity data for disulfiram across cancer cell lines were obtained from the Broad Institute PRISM Repurposing dataset integrated within the DepMap portal. PRISM screening measures pooled cell-line viability following exposure to repurposed compounds and reports drug-response metrics, including area under the dose-response curve (AUC). Lower AUC values indicate greater drug sensitivity. RNA expression profiles for targeted genes were integrated with PRISM drug-response data, and scatter plots were generated to evaluate the relationship between mRNA expression and disulfiram sensitivity across individual cell lines (n = 468). Cell lines were further stratified into low and high categories based on dependency or expression features to assess directional trends in drug responsiveness.

### LINCS-based identification of transcriptionally responsive compounds

To investigate transcriptional responses associated with Fanconi anemia (FA)-related marker genes, we analyzed chemical perturbation signatures from the LINCS L1000 Level 5 dataset (<u>GSE92742</u>, https://www.ncbi.nlm.nih.gov/geo/query/acc.cgi?acc=GSE92742), one of the largest publicly available resources describing compound-induced gene expression profiles)^17^. The Level 5 dataset contains consensus perturbation signatures generated using the Moderated Z-score (MODZ) algorithm, providing robust and reproducible estimates of transcriptional responses by integrating multiple biological replicates into a single consensus signature.

A predefined panel of 96 FA-associated marker genes contains 21 Candidate ALDH family genes involved in aldehyde detoxification, included based on their recurrent reduced expression in FA cells and their proposed role in protecting against endogenous aldehyde-induced DNA damage, and 75 differentially downregulated genes identified in FA-like cells, prioritized as potential rescue targets because their re-expression may improve cellular defects associated with the FA phenotype (**Table S1**). Expression values corresponding to the mapped marker genes were extracted from perturbation profiles generated across two predefined groups of DepMap cell lines representing low-(CRIPSR) dependency and high-(CRIPSR) dependency cellular phenotypes. The low-dependency group comprised 232 cell lines, whereas the high-dependency group comprised 84 cell lines (**Table S2**). Only chemical perturbation signatures (pert_type = trt_cp) derived from these cell lines were retained for downstream analyses. For every perturbation signature, normalized MODZ expression values of the mapped marker genes were extracted to generate compound-by-marker expression matrices. Replicate perturbation signatures corresponding to identical compound-cell line pairs (across different concentrations and treatment time points) were aggregated using the median to obtain unified transcriptional profiles. Four complementary expression matrices were subsequently generated, including a collapsed matrix containing all perturbation signatures, low- and high-dependency group-specific matrices, and a differential expression matrix representing the difference between the two cellular phenotypes (Low - High).

Before performing downstream analyses, the overall quality and statistical characteristics of the processed LINCS expression dataset were evaluated (**Supplementary Figure 7**). The distribution of normalized marker expression values was centered close to zero (**Supplementary Figure 7A**), consistent with the expected behavior of MODZ-normalized LINCS signatures and indicating successful normalization across perturbation profiles. Marker genes displayed varying levels of transcriptional variability, with a subset contributing disproportionately to overall transcriptomic heterogeneity (**Supplementary Figure 7B**). Gene-wise mean expression and sample-wise standard deviation analyses (**Supplementary Figure 7D**) demonstrated stable statistical characteristics without evidence of systematic bias, supporting the suitability of the processed expression matrices for downstream exploratory transcriptomic analyses.Downstream analyses included visualization of marker expression using heatmaps, t-SNE clustering to characterize global transcriptional variation, calculation of transcriptional response scores based on the euclidean norm of standardized marker gene expression profiles to identify highly responsive compounds, aggregation of compound-level scores across cell lines using the median, and correlation analysis among the highest-ranked compounds to identify compounds exhibiting similar transcriptional response patterns. An overview of the analytical workflow is presented in **Figure 1**, whereas a detailed description of the computational procedures is provided in **Supplementary Section S3**.

## RESULTS

### Fanconi anemia-like cellular models through functional dependency profiling

Despite the diverse tissue origins and cancer-specific genomic alterations present across the DepMap cell line collection, we hypothesized that dependency on the FA pathway could serve as a functional surrogate for an FA-like cellular state, enabling the identification of molecular determinants associated with FA vulnerability (**Figure 2A**). Rather than classifying cell lines according to tissue of origin or mutational status, we adopted an unbiased functional genomics approach in which cellular fitness following CRISPR-mediated gene disruption was used to define FA pathway dependency. This strategy allowed us to compare FA-like and control cell populations based on their reliance on the FA pathway rather than their intrinsic cancer characteristics.

To establish this framework, we first examined CRISPR gene dependency profiles for all 23 canonical FA genes across available 1178 cancer cell lines representing a broad range of tissue lineages in the DepMap database (**Figure 2B**). Several cell lines from the lymphoid lineage including OCILY19, SUDHL10, SUDHL8, and SUDHL5 exhibited the greatest overall dependency on the FA pathway, indicating a strong reliance on FA-mediated DNA repair for cellular fitness.

Because pathogenic variants in any one of the 23 FA genes can disrupt the FA pathway and give rise to the FA phenotype, we next derived a composite FA dependency score by calculating the median CRISPR dependency score of each 23 FA genes across all cell lines. This composite score was used to capture overall pathway dependency while minimizing gene-specific variability and technical noise across the dataset (**Figure 2C**). The overall median dependency score (0.26) across the complete FA gene set suggested that, as a pathway, FA genes exhibit low to moderate essentiality in the majority of cancer cell lines. Analysis of individual FA genes demonstrated that RAD51, RAD51C, and MAD2L2 were among the most universally essential genes across the DepMap cell line panel, consistent with their broader roles in homologous recombination and genome maintenance. In contrast, most FA genes exhibited context-dependent essentiality, with their disruption reducing cellular fitness rather than causing universal lethality (**Figure 2D**). This variability in pathway dependency indicates that only a subset of cell lines exhibit a marked reliance on the FA pathway, representing selective vulnerabilities that we refer to as FA-like cellular models. Based on the composite Chronos dependency score, cell lines were subsequently stratified into high, medium and low-dependency groups (**Figure 2E**). Because lower Chronos scores indicate higher dependency following gene disruption, this classification enabled the identification of FA-like cell populations and appropriate comparator groups for downstream analyses. We reasoned that comparing these dependency-defined groups would facilitate the discovery of molecular features associated with cellular fitness and reveal biological mechanisms that mimic key aspects of FA.

To further characterize these FA-like models, we integrated additional molecular features for each FA gene, including mutation status, CNA and mRNA expression across the same panel of cancer cell lines (**Figure 2F-G**; **Supplementary Figure 1**). These analyses revealed substantial molecular heterogeneity among FA genes across different tissue lineages. Although most FA genes exhibited relatively high mRNA expression levels (TPM), consistent with their fundamental roles in DNA repair and genome stability (**Figure 2F**), expression varied considerably among cell types (**Figure 2G**). Next, we examined the relationship between molecular alterations and functional dependency across the FA pathway (**Supplementary Figure 2**). As expected, CRISPR dependency and RNA interference (RNAi) score datasets showing the highest concordance (**Supplementary Figure 2A**). In contrast, mRNA expression patterns did not consistently correlate with gene essentiality, as many FA genes remained highly expressed in cell lines that showed little dependency on them for survival (**Supplementary Figure 2B**). These observations also suggest that transcriptional abundance alone is an insufficient predictor of FA pathway dependency and that compensatory or redundant mechanisms likely buffer the functional consequences of FA gene perturbation. Importantly, this discordance provides an opportunity to identify the molecular determinants and compensatory pathways that underlie FA-like cellular vulnerability.

### Molecular features associated with CRISPR dependency in FA-like phenotypes

To explore key molecular determinants associated with FA-like phenotypes using cell lines, we employed a combination of supervised learning and statistical modeling. We trained Random Forest (RF), XGboost (XGB) classifier and soft voting ensemble on multi-omic features, mRNA expression, CNV, and mutation status, to predict gene dependency categories (low, medium and high). This method enables both predictive modeling and variable importance ranking, making it well-suited for the integration and interpretation of high-dimensional datasets. We also applied SHAP analysis of the XGBoost model to identify features contributing most strongly to dependency classification. Additionally, t-tests were used to assess statistically significant differences in feature distributions between cell lines with high and low dependencies on specific FA genes.

Interestingly both RF and XGB classifiers gave consistent output that made it confident, and their combined soft-voting ensemble discriminatory score was found to be modest. The models are reasonably good at identifying “High” CRISPR dependency but often misses actual cases (∼50% precision and ∼50% recall) (**Supplementary Figure 3A-C**). Low Class also showed similar performance with lower precision and recall. “Medium” Class is poorly predicted with lowest precision (<40%), and recall (<35%), indicating the model often guesses Medium when it shouldn’t (**Supplementary Table 1**). This implies the robust comparison group would be to analyze the low and high dependency features, while median leave out of analysis to see the clear differences in two extremes. Hence we opt to do further analysis using two distinct cell groups. The models are getting better at identifying both high and low Class CRISPR dependency (>60% precision and recall) (**Figure 3A-C**). Furthermore, ROC analysis demonstrated that the models exhibited stronger discriminative ability, with AUC values substantially above the random baseline (AUC ∼ 0.7) with low vs high comparison (**Figure 3D**). Model performances were also assessed using confusion matrices and finding moving from binary (low vs high, **Figure 3E-F**) to multiclass classification (low vs medium vs high, **Supplementary Figure 3E-F**) reduces per-class accuracy, as expected. Errors concentrate toward the medium group, further emphasizing that it represents a biologically continuous transition zone rather than a discrete group (**Supplementary Figure 3D-F**).

Each machine learning approach, including RF, XGB, and SHAP generated a ranked list of molecular features capable of discriminating FA dependency groups using both binary (Low vs. High) (**Figure 3G-I)** and multiclass (Low, Medium, and High) classification models**; Supplementary Figure 3G-I)**. Collectively, these analyses identified potential candidate molecular determinants associated with FA-like cellular states (**Supplementary Table 1**). To complement this analysis, we performed differential feature analysis using Student’s *t*-test to quantify molecular differences between the low- and high-dependency groups. Although the predictive performance and feature importance scores were generally modest, likely reflecting the intrinsic heterogeneity of large-scale multi-omic datasets and the complex molecular landscape of cancer cell lines, the models consistently identified informative features that distinguished dependency groups. The feature sets comprised multiple molecular data types, including somatic mutations, copy number variations (CNVs), and mRNA expression profiles. Notably, mRNA expression features predominated among the highest-ranked predictors, suggesting that transcriptional programs contribute more substantially to FA pathway dependency than genomic alterations alone (**Supplementary Table 1**). While the top-ranked genes were not directly linked to canonical FA biology, they may represent previously unrecognized regulators, compensatory pathways, or downstream effectors associated with FA-like cellular vulnerability. These candidate genes were therefore subjected to further integrative analysis in subsequent sections to evaluate their biological relevance. Comparison of the prioritized gene sets revealed only limited overlap between RF and the XGBoost/SHAP models (**Supplementary Figure 3J, Figure 3J**). This discrepancy is expected, as RF preferentially identifies robust features with stable, averaged effects across the dataset, whereas XGB and SHAP are more sensitive to conditional, nonlinear, and interaction-dependent relationships.

### mRNA expression profiles exhibited consistent predictive signatures matching with Fanconi Anemia patient samples

We next performed differential analysis between the low- and high-dependency groups using an unpaired t-test across all molecular features (**Supplementary Figure 4A**). This analysis enabled us to quantify the magnitude of differences between the two groups by calculating fold changes for each molecular feature. Among the molecular data types examined, mRNA expression exhibited the greatest quantitative differences, with substantially larger fold changes than mutation, copy number variation, methylation, or CRISPR dependency data, suggesting that transcriptional alterations most strongly distinguish the two dependency groups (**Figure 4A**). This observation is consistent with RF, XGB analysis, indicating the predictability of CRISPR dependency in those cell lines are largely influenced by the mRNA expression. However, CNV and mutation were called sizable representations from the RF and XGB analysis (**Supplementary table 1-2**). We hypothesize that, compared with DNA-based alterations (mutations, CNVs, and LOH), mRNA expression provides a more proximal representation of cellular phenotype because it integrates the cumulative effects of genetic, epigenetic, and environmental regulation. As the functional output of upstream molecular events, transcriptional profiles capture dynamic pathway activity and cellular state, whereas mutations and CNVs may reflect historical tumor evolution, clonal selection, or long-term culture adaptation and are not always directly relevant to the biological processes underlying inherited disorders such as FA^22,23^.

The top-ranked with high differential expressed genes mapped to biological processes that potentially modulate FA phenotypes, while canonical FA genes were not among the top, consistent with outcome from classification based predicting features. To investigate coordinated biological processes rather than individual gene effects, we performed pathway-level analyses using Gene Set Variation Analysis (GSVA) and single-sample Gene Set Enrichment Analysis (ssGSEA). These approaches estimate pathway activity scores for each sample, enabling comparison of coordinated pathway activity between the low- and high-dependency groups (**Figure 4B; Supplementary Figure 4B**).

Several significantly enriched pathways were consistent with previously reported metabolic and cell-cycle alterations associated with FA. Specifically, estrogen response, xenobiotic metabolism, cholesterol/fatty acid metabolism, and androgen response pathways were significantly downregulated in the high-dependency group (low CRISPR score) (ΔGSVA ≈ 0.03 to 0.05; FDR q < 10^-5^). Conversely, pathways related to E2F targets, the G2/M checkpoint, and mTORC1 signaling were significantly enriched (ΔGSVA ≈ −0.04; FDR q < 10^-5^), indicating increased proliferative and cell-cycle activity. However, the increased proliferative activity likely reflects the inherent cancerous nature of these cell lines. In addition, glycolysis, UV-response, and p53-associated pathways exhibited modest but statistically significant differential enrichment. These observations are consistent with previous studies reporting metabolic reprogramming, oxidative stress responses, and endocrine dysregulation as FA phenotype ^24,25^, although the direction and magnitude of these changes are known to be cell-type dependent.

In order to exclude the chances of cell line (lineage) specific impartiality, we compared the outcome with curated genes from FA patients’ samples. For this, we reanalyze the gene expression across independent transcriptomic datasets containing normal or healthy vs FA-patient derived samples including GEO microarray ^18^, bulk RNA-sequencing ^19^, and single-cell RNA-sequencing ^20^ (**Supplementary Fig 4C-E**). Details are provided in the supplementary material. We applied directional concordant analysis to match the genes from the FA patients sample compared the direction and magnitude of differential expression between the DepMap CRISPR-derived gene prioritization and three transcriptomic datasets (**Supplementary Figure 4F-H**). Importantly, our analysis revealed substantial shared genes that exhibited the same direction of regulation across independent datasets (**Figure 4C-D)**. These findings indicate that while the magnitude of gene expression changes varies between platforms, the overall direction of transcriptional regulation is preserved for a substantial proportion of genes, supporting the robustness of the identified disease-associated transcriptional signatures. For prioritization, we primarily relied on the bone marrow microarray dataset reported by Vanderwerf *et al.* 61 genes found to be directionally matched with either low (42 genes) or high (19 genes) expression with the curated gene list of predictive models and FA patients (**Figure 4E**).

Enrichment analysis of low expressed genes were interpreted using top-ranked terms from relevant databases, including gene ontology, molecular function, and Reactome to identify pathway-level biological themes (**Figure 4F-G**). Notably, FA partnering pathways were particularly enriched including ethanol oxidation, retinoic acid synthesis, aldehyde metabolism and neutrophil degranulation. Interestingly, several genes within the developmental biology domain were highlighted via Orphanet augmented (**Supplementary Figure 4I**). It has been found to be altered in FA-like cells that are also key players in broader congenital/ developmental disease contexts, suggesting that the molecular networks disrupted in FA, regardless of cellular context, overlap with those that govern embryonic morphogenesis and organ development.

### Hematopoietic lineage displays characteristic and predictable traits

Next, we investigated whether lineage-specific vulnerabilities associated with FA could contribute to the BMF phenotype. To address this, we integrated CRISPR dependency and gene expression data across all cell lines and performed principal component analysis (PCA) to evaluate lineage-specific patterns. Interestingly, cell lines are predominantly clustered according to their tissue of origin rather than their dependency profiles. For example, myeloid and lymphoid cell lines largely grouped together and showed limited separation based solely on CRISPR dependency, suggesting that additional molecular features or further stratification may be required to better resolve FA-like phenotypes within hematopoietic malignancies (**Figure 5A**). Consistent with these findings, hematopoietic cell lineages, comprising both myeloid and lymphoid cells, were the most discriminative lineage in the dependency analysis, and were enriched for the high-dependency group **(Figure 5B**). This observation is consistent with the well-established hematopoietic vulnerability of FA, in which defects in DNA repair preferentially affect hematopoietic stem and progenitor cells, ultimately leading to bone marrow failure. Lymphoid lineage in particular clusters together with highly sensitized cells towards FA genes as shown in UMAP analysis that preserves non-linear local neighborhood structure, allowing coordinated lineage-specific transcriptional programs to emerge as distinct clusters (**Supplementary Figure 5A**). The observed clustering demonstrates that the prioritized gene signature preserves biologically meaningful transcriptional organization across diverse cancer cell types while maintaining associations with FA dependency. The hierarchical cluster of gene expression also makes a cluster of lymphoid in a larger group, however there’s a mixed trend of lower and high CRISPR dependency groups (**Figure 5C**). It suggests the selective sensitivity of individual genes is unlikely to make any significant impact in the larger extent instead of a compound effect of multiple gene action. Top ranked differential expressed genes included CD70 (immune activation), GYPC (erythroid integrity), ADGRG6 (cell-matrix signaling), and NLRP1 (inflammasome activation), highlighting coordinated immune, hematopoietic, and stress-response alterations (**Supplementary Figure 5B**). SHAP interaction analysis captures modest biologically meaningful gene-gene dependencies, approximating functional epistasis and synthetic lethality/viable patterns. This suggested genetically similar cell lines may occasionally exhibit divergent SHAP interaction scores, and context-dependent pathway wiring rather than transcriptomic similarity alone (**Supplementary Figure 5C**).

**Figure 5:**
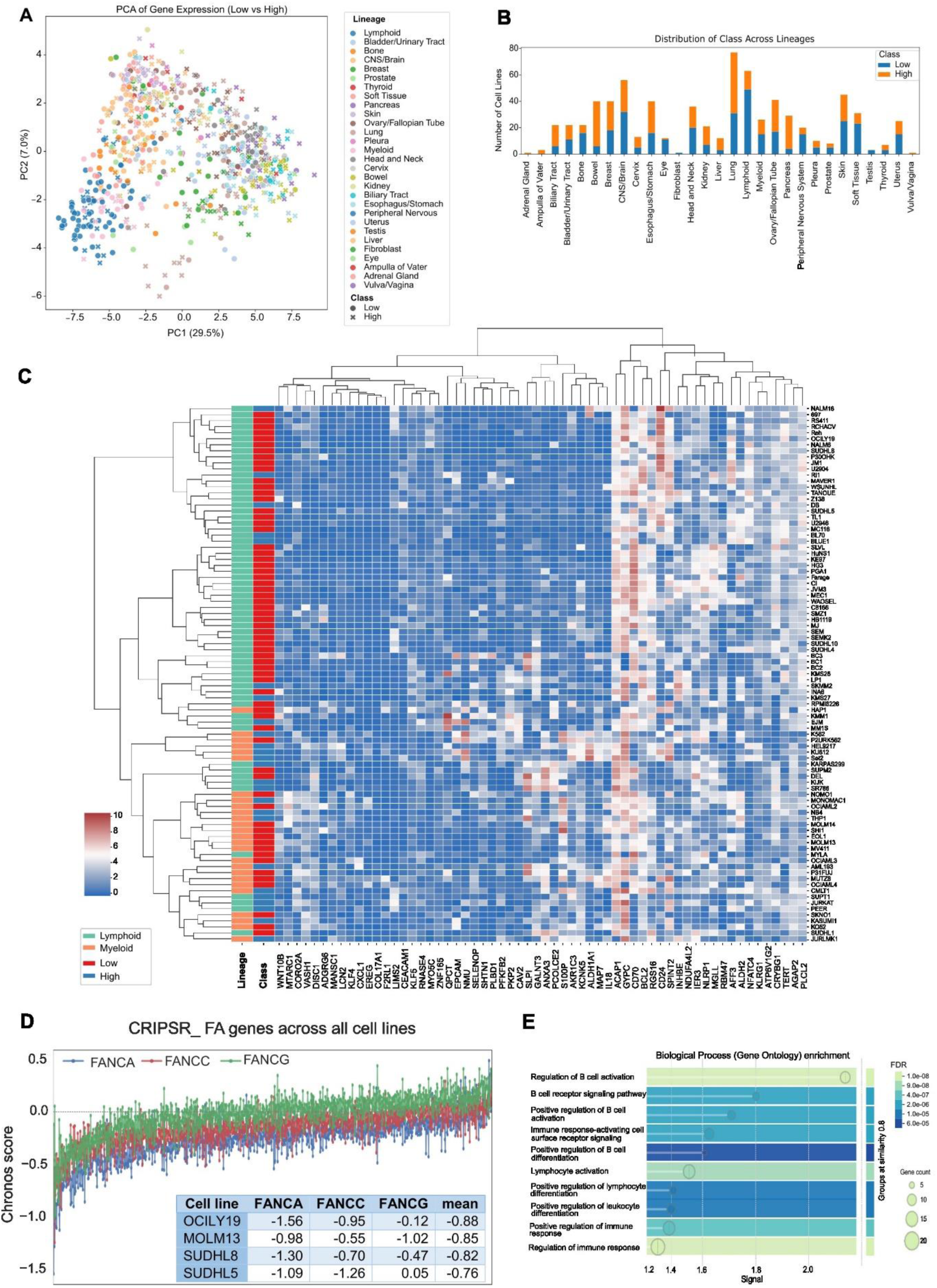
Hematopoietic cell lineages exhibit distinct molecular characteristics. **A)** PCA analysis displayed category and class discrimination among cell lines, highlighted hematopoietic lineage as the most discriminatory, and exhibited higher CRISPR dependencies. Samples are colored according to tissue lineage and shaped according to FA dependency class (Low or High). **B)** Distribution of cell lines based on their lineages and categories with CRISPR dependency group (low/high). **C)** Heatmap of 61 concordant mRNA expression annotated with the CRISPR dependencies groups across lymphoid and myeloid cell line. **D)** Cell lines were ranked by mean FA dependency scores of FANCA, FANCC, and FANCG, and the dependent cell lines were highlighted by lineage. Lymphoid cells predominantly represent the most vulnerable subset with lowest Chronos scores when compared to the top FA causing genes vs rest of the genes across all cell lines. **E)** Top 50 differential RNA of the four FA-dependent cell lines were subjected to enrichment networks using https://string-db.org/ and presented GO biological process with the high confidence threshold.

Next, we identified the most vulnerable subset of cell lines using the major FA-causing genes FANCA, FANCC, and FANCG, which together account for approximately 80-90% of clinically diagnosed FA cases ^26^. The combined FA dependency score revealed a subset of cell lines with markedly lower Chronos values for FANCA, FANCC, and FANCG, indicating increased reliance on the FA pathway (**Figure 5D, Supplementary Figure 5D**). The strongest dependent lines were enriched for specific lineages, including lymphoid (e.g., OCILY19, SUDHL8, SUDHL5) and myeloid (MOLM13) cell lines. Comparison of RNA profiles between the top FA-dependent cell lines and the remaining lines showed differential expression across multiple genes (**Supplementary Figure 5E**), supporting a distinct transcriptional state associated primarily with hematopoietic and lymphocyte-related biological processes and enrichment networks (**Figure 5E, Supplementary Figure 5F**).

### Cross-talk between endogenous aldehyde detoxification and the Fanconi anemia pathway

To investigate the biological basis of FA-like dependency patterns across cell lines, we examined the functional relevance of the identified genes and pathways. Pathway enrichment analyses consistently highlighted aldehyde metabolism and closely related metabolic processes as significantly enriched across multiple independent analyses (**Figure 4F-G, Supplementary Figure 6A-B**), suggesting a recurrent biological theme. Consistent with these pathway-level observations, several key genes involved in aldehyde detoxification, including ALDH2, ALDH1A1, ALDH1A3, and AKR1C3, were among the most significantly downregulated genes in the high-dependency group (**Figure 5A)**, indicating that impaired aldehyde metabolism may contribute to the reduced dependence on the FA pathway. Interestingly, the genes emerging from our predictive modeling converge strongly on metabolic buffering of aldehyde and lipid-derived electrophiles (**Figure 5B**). ALDH1A1 and ALDH2 catalyze detoxification of retinaldehyde and acetaldehyde, respectively, preventing accumulation of aldehyde adducts that overwhelm DNA repair machinery ^9,28^. Likewise, AKR1C3 may contribute directly to the detoxification of reactive carbonyl and aldehyde species that accumulate during oxidative stress, whereas MGLL may indirectly influence aldehyde burden by regulating lipid metabolism and the formation of lipid peroxidation-derived aldehydes ^29,30^ (Fig.5 B).

Interestingly, the genetic dependencies of aldehyde metabolism genes are minimal with substantial expression of several of them, and have poor correlation across the cell line panel (**Figure 5C**). However, reduced expression of these enzymes was modestly associated with the high-dependency cell lines (**Figure 5D**), while not particularly enriched across all hematopoietic lineages (**Figure 5E**), suggesting a model in which diminished intrinsic aldehyde detoxification capacity functionally synergizes with FA-like defects to increase reliance on compensatory survival programs (**Supplementary Figure 6C-E**). Four lymphoid cell lines, OCILY-119, SUDHL-5, SUDHL-8 and SUDHL-10 were particularly lower to the most prominent FANCA gene dependency, that expressed no or very little ALDH2 and partner genes further supported the physiological connection of aldehyde detoxifying pathway in FA context **(Supplementary Figure 6F**). Notably, the coordinated downregulation of multiple aldehyde-handling pathways rather than a single enzyme implies broader metabolic vulnerability rather than isolated pathway perturbation.

### Transcriptional rescue scoring identifies candidate compounds for FA-phenotype

Next, we sought to identify small molecules capable of rescuing the FA-like phenotype by restoring the expression of low-expressed genes in FA-like cells, using the LINCS L1000 Level 5 dataset (GSE92742). **Figure 7** summarizes the transcriptomic characterization of LINCS perturbation signatures using a panel of FA-associated marker genes across two predefined groups of DepMap cell lines representing low- and high-dependency phenotypes. More details on the analytical workflow are provided in **Supplementary Section S3**.

**Figure 6.**
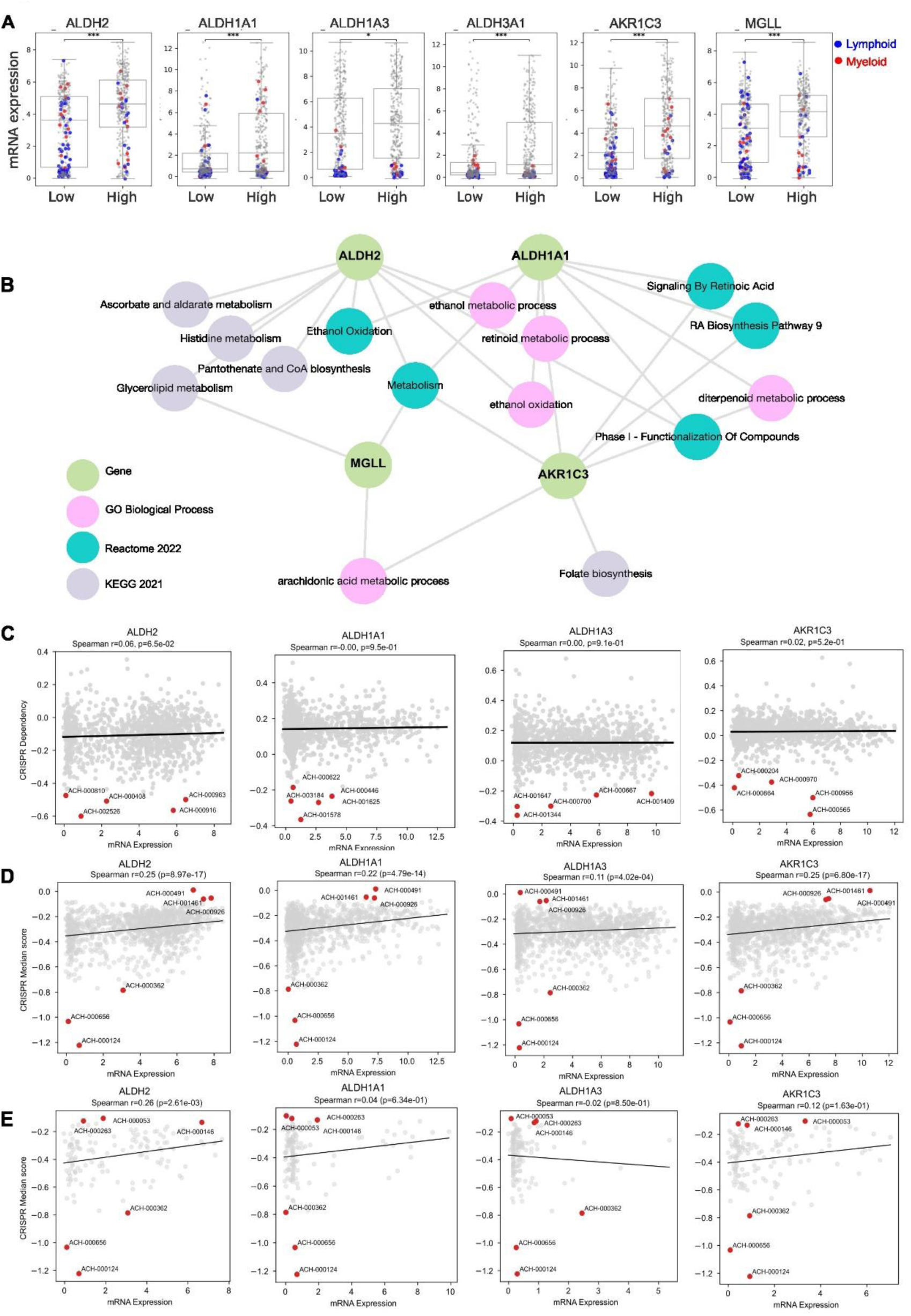
Endogenous aldehydes and Fanconi Anaemia pathway interactions. **A)** Boxplot highlighted differential expression of top aldehyde dehydrogenase family members across all cell lines. Colors are indicating respective lymphoid/myeloid lineages within the plot. Statistical significance was determined using t-test. Significance levels are denoted as P < 0.05 (*), P < 0.01 (**), and P < 0.001 (***); ns = not significant. **B)** Key genes were submitted to web-based gene-set enrichment analysis platform Enrichr (Ma’ayan Lab) to visualize the interaction across multiple functional libraries (biological process, Reactome and KEGG). **C)** Scatter plot showed member genes and their correlation with expression and CRISPR dependency scores. **D-E)** Individual members and their expression were plotted against all stratified cell line’s median CRISPR score **D)** across all cell lines. **E)** only in hematopoietic cell lines. Top 3 (high and low dependency score) cell lines were highlighted in red color.

**Figure 7:**
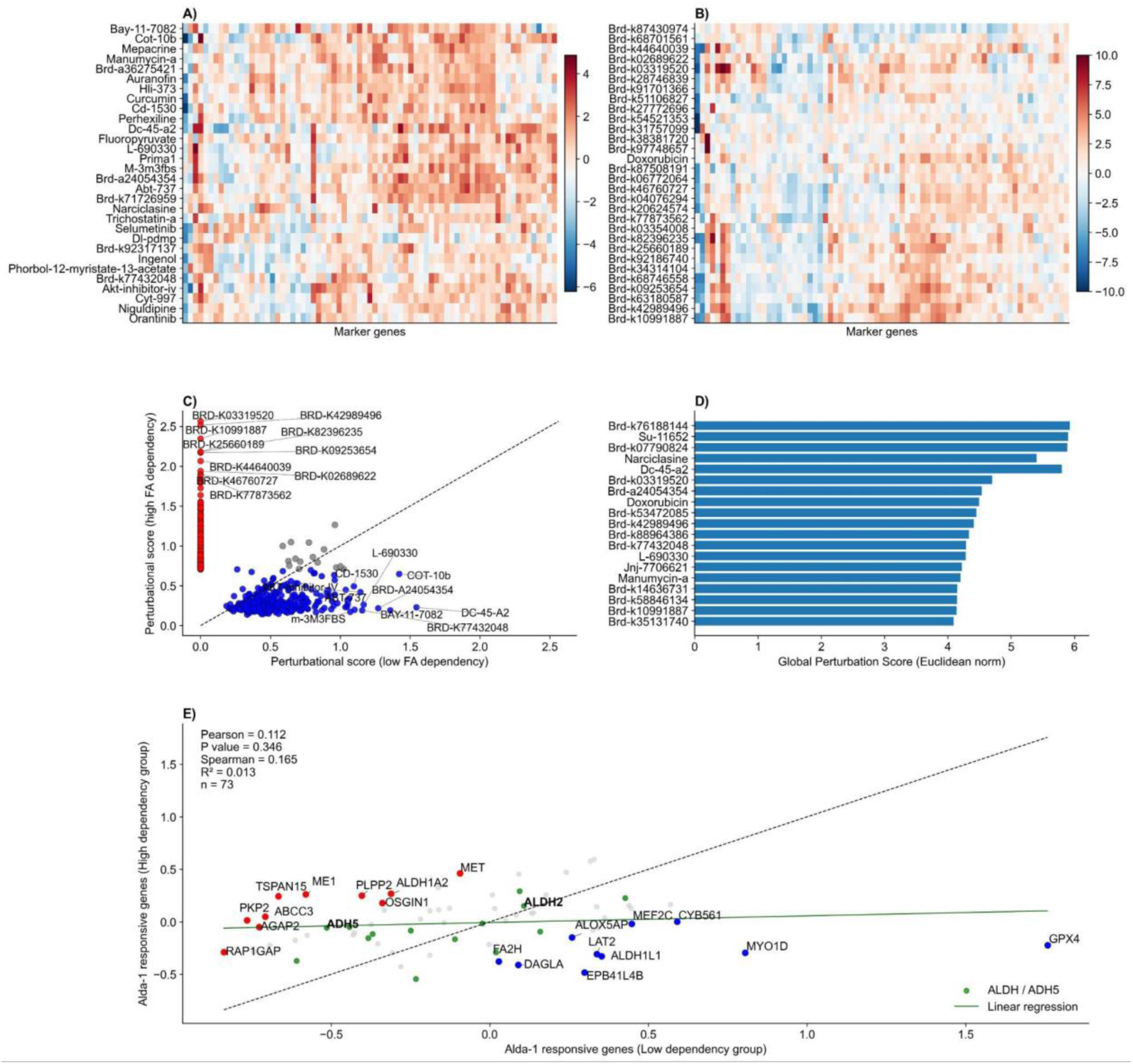
Global transcriptional characterization of LINCS marker expression profiles across compounds. **(A-B)** Heatmap showing the 30 most transcriptionally variable compounds within **(A)** the low-dependency and **(B)** high-dependency cell line group based on marker gene expression. Compounds and marker genes were hierarchically clustered based on their expression profiles, with dendrograms omitted for visualization. Heatmap colors represent standardized expression (z-score), where red and blue indicate relatively increased and decreased expression, respectively. **(C)** Scatter plot comparing compound perturbational activity scores between the low- and high-dependency groups. Perturbational activity scores were calculated as the mean absolute MODZ-normalized marker gene expression across all mapped FA-associated genes, and duplicate compound signatures were collapsed by retaining the highest-scoring instance for each compound. The dashed diagonal indicates equal perturbational activity between the two cellular phenotypes, while labeled compounds represent the strongest perturbagens. **(D)** Top 20 compounds ranked by transcriptional response score, calculated as the Euclidean norm of gene-wise standardized marker gene expression values, highlighting compounds producing the strongest global transcriptional perturbations. **(E)** Scatter plot showing the relationship between Alda-1-induced transcriptional responses of 73 marker genes in the low- and high-dependency groups, with each point representing an individual gene. Red and blue points denote the 10 genes showing the largest positive and negative differences between the groups, respectively, while orange points indicate aldehyde dehydrogenase-related genes, including highlighted ALDH2 and ADH5. The dashed black line represents the identity line (y = x), while the solid regression line represents the fitted linear relationship between gene responses in the two groups. Pearson and Spearman correlation coefficients and their corresponding P-values quantify linear and rank-based concordance, respectively, while R² indicates the proportion of variation explained by the linear regression (n = 73 genes).

The processed expression matrices revealed substantial heterogeneity in compound-induced transcriptional responses across the mapped FA-associated marker genes (**Figure 7A-B**). Although many compounds produced relatively modest transcriptional changes, distinct expression patterns were evident across both low- and high-dependent cellular phenotypes. Hierarchical clustering of both compounds and marker genes further revealed distinct groups of perturbagens with coordinated transcriptional response patterns within each dependency phenotype.

Comparison of perturbational activity scores between the low- and high-dependency groups demonstrated limited concordance across perturbagens, indicating that compounds eliciting strong transcriptional perturbations in one phenotype did not necessarily produce responses of comparable magnitude in the other (**Figure 7C**). Compound-level response scores were subsequently ranked to identify the most transcriptionally responsive compounds (**Figure 7D**). Several compounds exhibited substantially larger response scores than the remainder of the dataset, indicating their ability to induce coordinated transcriptional perturbations across multiple FA-associated marker genes. These highly responsive compounds represent promising candidates for subsequent mechanistic and functional investigation.

We selected Alda-1, a prototypic ALDH2 activator, for a focused mechanistic case-study analysis. Lower ALDH2 expression is a major feature of the FA-like phenotype (high-dependency group) and has also been observed in FA patients (Figure 7E). Although Alda-1 produced only a modest increase in ALDH2 expression in the high-dependency group, it increased the expression of several other genes that were reduced in the high-dependency group, including aldehyde-metabolism-associated genes such as ADH5, which can provide compensatory formaldehyde-detoxification activity when ALDH2 function is compromised (**Figure 7E**). These findings suggest that the transcriptional response to Alda-1 may extend beyond direct modulation of ALDH2 and involve a broader aldehyde-detoxification response in the FA-like cellular context.

Together, these observations indicate that chemical perturbations induce diverse transcriptional programs across the FA-associated marker panel and that the magnitude and direction of these responses differ considerably between the predefined cellular phenotypes. The Alda-1 case study further illustrates how the framework can move from global compound prioritization to gene-level interrogation of a selected perturbagen, identifying transcriptional responses that may contribute to restoration of pathways dysregulated in the FA-like phenotype. Thus, the analyses provide a comprehensive transcriptomic characterization of compound-induced responses across FA-associated marker genes and establish a robust framework for prioritizing transcriptionally responsive compounds for further biological validation.

### Drug affinity mapping prioritizes aldehyde detoxification enzymes as pharmacologically tractable nodes

Next we sought to explore if any pharmacological intervention is able to rescue the endogenous cellular stress via suppression of higher expressed genes and vice versa in FA-like cells. To this end, we characterized pharmacological interactions involving prioritized lists of targets by programmatically mapped gene symbols for 61 genes to their corresponding human UniProt identifiers, consistent with ChEMBL’s target annotation framework. First, we curated the differential expressed genes and put them into the drug-gene interaction pipeline to discover the binding affinities to the small molecule. Our drug-target affinity analysis identified 12 approved drugs with high-confidence binding to at least one of the molecular candidates prioritized in our model, thus demonstrating putative engagement (**Figure 8A**). Notably, disulfiram emerged as a multi-target modulator of lipid and aldehyde-metabolism pathways, with predicted high-affinity binding to ALDH1A1, ALDH2, and MGLL (MAGL). Experimental literature supports disulfiram as an inhibitor of human purified MAGL ^31^, whereas ALDH2 and and ALDH1 isoforms are the classical canonical aldehyde-detoxifying target of disulfiram ^32^., although MGLL is not a canonical aldehyde dehydrogenase, its inhibition may indirectly influence reactive carbonyl stress through altered lipid turnover and endocannabinoid metabolism.

**Figure 8.**
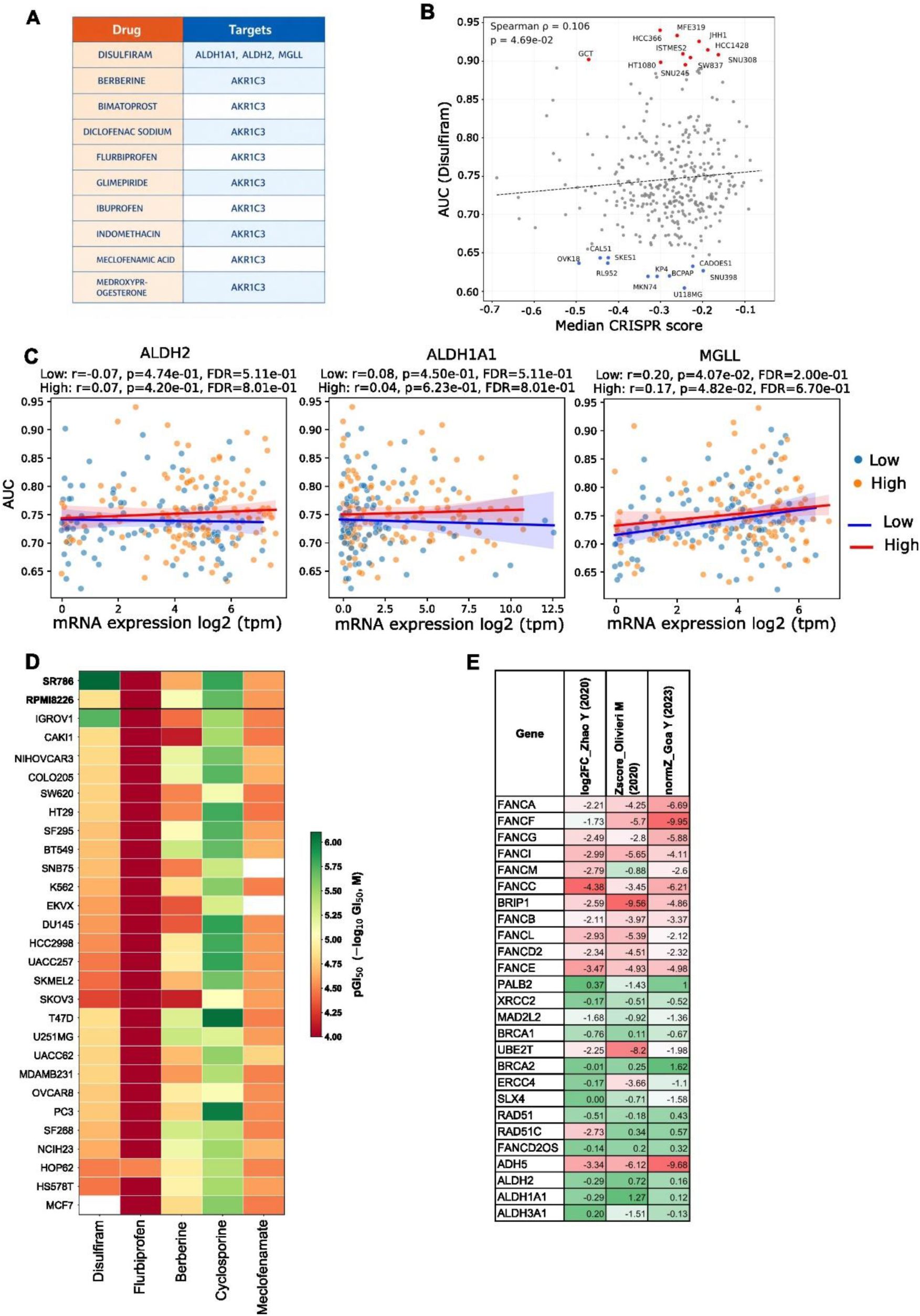
Drug affinity mapping identifies aldehyde detoxification pathway as therapeutic targets. **A)** Approved drugs targeting candidate genes based on drug-target affinity data from ChEMBL (version 36) with nine small molecules to be interactive with AKR1C3, including berberine, bimatoprost, diclofenac sodium, flurbiprofen, glimepiride, ibuprofen, meclofenamic acid, medroxyprogestron and indomethacin. **(B)** Scatter plot showing the correlation between median dependency scores and disulfiram AUC values across common cell lines. Each point represents one cell line, and the dashed line represents the linear regression fit. **(C)** Scatter plots showing mRNA expression versus drug response (AUC) across DepMap cell lines, stratified by CRISPR dependency status (low and high dependency). Each point represents an individual cell line. Spearman correlation coefficients and corresponding *P* values were calculated separately for each group. **(D)** Heatmap showing pGI50 values for 29 cell lines including 2 Lymphoid cells highlighted in bold, treated with top 5 drugs. Higher pGI50 values (green) indicate greater drug sensitivity, whereas lower values (red) indicate relative resistance **(E)** Knockout effect scores of FA and aldehyde metabolism genes from three independent CRISPR knockout studies evaluating formaldehyde sensitivity in RPE1 and K562 cell lines, where knockout of genes with negative Z-scores/ LFCs confers sensitivity to formaldehyde.

To determine whether the predicted drug-target interactions translated into phenotypic responses, we interrogated disulfiram sensitivity using the PRISM Repurposing dataset from DepMap. Overall, disulfiram exhibited limited anticancer activity, with relatively high AUC values across most cell lines **(Figure 8B)**. A modest association was observed between FA-related dependency scores and disulfiram AUC, with higher drug efficacy largely observed among low-dependent cell lines. In contrast, several highly dependent cell lines exhibited markedly lower AUC values, indicating increased sensitivity to disulfiram, and providing phenotypic support for the predicted vulnerability. We further stratified cell lines according to low and high FA dependency and evaluated disulfiram sensitivity in relation to the expression of its predicted targets (ALDH2, ALDH1A1, and MGLL) **(Figure 8C)**. No significant association was observed between target expression and drug response, suggesting that expression of these enzymes alone is insufficient to predict selective sensitivity to disulfiram. To further evaluate this observation, we examined publicly available NCI-60 drug-response data for the prioritized approved compounds **(Figure 8D)**. Among the available datasets, disulfiram demonstrated overall weaker anti-proliferative activity as a single agent, with peak sensitivities observed in lymphoid cell line SR786 (pGI50 = 6.0) and ovarian cancer cell lines IGROV1 (pGI50 = 5.73). Interestingly, SR786 belongs to the high-dependency group and concurrently exhibits low mRNA expression across several aldehyde dehydrogenase genes (ALDH2, ALDH1A1, ALDH1A3, and ALDH3A1). This observation mechanistically corroborates our hypothesis that FA-derived malignancies, with compromised ALDH detoxification activity can be effectively targeted by disulfiram. Recent functional genomic studies provide additional biological context for these findings. Genome-wide CRISPR screens performed under formaldehyde stress identified multiple components of the FA pathway together with ADH5 as critical determinants of formaldehyde tolerance in both hTERT-immortalized RPE1 cells and K562 leukemia cells ^33–35^ **(Figure 8E)**. Notably, ADH5 emerged as the principal aldehyde-detoxification enzyme whose loss consistently sensitized cells to endogenous formaldehyde, reinforcing the functional link between aldehyde metabolism and FA pathway integrity. Collectively, these findings suggest that although inhibition of aldehyde-metabolizing enzymes alone does not produce selective cytotoxicity in cancer cell lines, impaired aldehyde detoxification remains a central determinant of cellular vulnerability.

Interestingly, our analysis identified nine small molecules to be interactive with AKR1C3 (**Figure 8A**). This suggests that AKR1C3 may represent a broader pharmacologically tractable node within the aldehyde-metabolism network, with several chemically diverse compounds converging on the same target. In addition, we included cyclosporine, which targets ABCC3, a gene encoding an ATP-dependent efflux transporter. ABCC3 consistently emerged as a predictive marker across multiple analyses, with lower mRNA expression observed in high-dependency cell lines. It also demonstrated increased sensitivity across the cell line panel. Overall, our analysis predicted drug-target interactions primarily involved enzymes that were already transcriptionally reduced, including enzymes involved in endogenous aldehyde detoxification and redox homeostasis. Therefore, while these compounds are unlikely to reverse the molecular phenotype associated with FA deficiency, they may represent opportunities to exploit existing metabolic vulnerabilities in FA-dependent or FA-transformed cancers through a synthetic-lethal mechanism.

## DISCUSSION

This study addresses the limited availability of robust FA disease models by leveraging the systematic integration of multi-omics datasets using conventional cell line data. We highlighted key molecular vulnerabilities and predictive biomarkers through multi-omics integration in a novel and comprehensive manner not previously undertaken for FA. Since only few non-cancerous in vitro models exist that reliably recapitulate FA biology ^13^. Even iPSC-derived models or transformed cancer cell lines derived from FA patients are limited in number and accessibility ^12,14^. Thus, we stratified cancer cell lines that exhibit features mimicking the FA phenotype, specifically, those whose cellular fitness is uniquely sensitive to the loss of FA-associated genes, effectively mimicking key features of FA-related vulnerabilities (**Figure 2E**). These vulnerable cell lines were identified with context-specific dependencies, and served as functional surrogates to study FA-associated deficiencies. Our analysis revealed transcriptomic data offer a more immediate and integrative proxy for cellular phenotype, also in comparison with FA patients derived transcriptomics data. Since expression changes often reflect the downstream consequences of genomic events, thereby providing greater insight into functional biological modules ^22,23^. In contrast, mutations and CNVs in cancer-derived cell lines may largely represent the legacy of tumor evolution and may not be directly relevant to FA biology.

Integrative analysis also revealed genes involved in diverse functional processes, supporting our hypothesis that predictive biomarkers are more likely to emerge from compensatory pathways and gene activities rather than solely from canonical FA pathway genes (**Figure 3-4**). This is consistent with the view that FA is a multisystem disorder with clinically significant metabolic and endocrine abnormalities, including disturbances in lipid metabolism ^26^. Some of the identified genes belong to biological programs known to be imbalanced in FA patients and in broader BMF, including pathways related to cholesterol and fatty acid metabolism. In parallel, the lower androgen-response signature observed in FA-like cells may be relevant to the clinical utility of androgens in FA-associated BMFS, where androgens remain a recognized supportive treatment option ^36^. The outcome also nominated key players whose involvement extends beyond FA into broader congenital disease contexts (**Figure 4F**), which is in line with the well-established fact that FA is characterized by congenital malformations and multi-organ developmental defects ^37^. However, the ability to distinguish these phenotypes within cell lines is notable and highlights the intrinsic, context-dependent behavior of cellular models.

Our integrative analysis identified multiple genes involved in aldehyde metabolism and associated detoxification pathways as key discriminative features between FA-like and non-FA cellular states (**Figures 5-6**). Notably, these enzymes were transcriptionally downregulated, suggesting impaired detoxification capacity in the FA-like state. ALDH2 has been shown as a critical factor in clearing acetaldehyde-induced damage that FA-deficient cells cannot repair ^9^. These aldehyde-metabolizing enzymes play a compensatory protective role in cells where the FA DNA repair pathway is defective. Here this is also to consider that the cancer cell lines with FA gene dependencies are enhanced with lower ALDH enzymes suggesting a synthetic lethal combination against FA transformed cancers. Clinical evidence supports this mechanism, in Japanese FA patients, the dominant-negative ALDH2 variant was associated with earlier BMF and more congenital anomalies, indicating that reduced aldehyde clearance worsens disease severity ^38^. These observations highlight aldehyde metabolism as both a mechanistic contributor to FA pathogenesis and a potential source of biomarkers capable of distinguishing FA-related cellular states.

The biological origins and evolutionary drivers of cancers arising in FA remain poorly defined, and there are currently no proven strategies to prevent or delay malignant transformation in this population. Furthermore, most individuals with FA cannot safely receive standard DNA-damaging therapies, such as DNA cross-linking chemotherapeutics or ionizing radiation, because of their inherited hypersensitivity to genomic injury ^26^. Within this context, our pharmacologic profile conceptually aligns with current models in FA biology in which disulfiram’s engagement of multiple aldehyde-buffering enzymes suggests that, while it may be mechanistically attractive as a synthetic-lethal sensitizer in FA-deficient malignancies, it is unlikely to be protective in FA-like contexts where aldehyde detoxification capacity is already functionally attenuated. In contrast, our iLINCS analysis provides a complementary approach for identifying pharmacological strategies aimed at the FA-like cellular state itself. By identifying compounds whose perturbational transcriptional profiles oppose the FA-associated expression signature, this analysis highlights candidate compounds that may have the potential to modulate or partially reverse molecular features associated with the FA-like phenotype. Although transcriptional reversal does not by itself establish therapeutic efficacy, this approach provides a starting point for prioritizing pharmacological interventions that could be explored further in FA models. Activation of aldehyde-detoxifying pathways may represent a more rational protective strategy for FA and FA-like disease states, where endogenous aldehydes amplify replication stress and DNA crosslink burden. Alda-1, the prototypic ALDH2 activator, directly rescues ALDH2 function and has shown cytoprotective effects in multiple preclinical injury models, supporting the concept that ALDH2 activation lowers toxic aldehyde burden and preserves cellular integrity ^39^. The translational relevance of this approach is further strengthened by the development of next-generation ALDH2 activators such as AD-9308, a potent and water-soluble compound that improved metabolic and cardiac phenotypes in preclinical models ^40^. Importantly, this biology has now moved into clinical testing: FP-045 (mirivadelgat), an ALDH2 activator, has been registered in a phase 1/2 dose-escalation study in patients with FA to assess safety, tolerability, pharmacokinetics, and preliminary biological activity (https://clinicaltrials.gov/study/NCT04522375). In cancer models, ALDH2 modulation has also shown context-dependent effects; for example, Alda-1 restored ALDH2-mediated alcohol metabolism and suppressed NF-κB/VEGFC signaling in head and neck cancer cells, reinforcing the idea that aldehyde detoxification may be protective in inherited FA but therapeutically exploitable in FA-associated malignancies ^41,42^

This study has several important limitations that should be acknowledged. First, our findings are derived primarily from in silico analyses of publicly available multi-omics and CRISPR dependency datasets, which despite their depth and breadth, may not fully capture the physiological complexity of primary FA tissues. However, rigorous multi-layered validation using RNA-sequencing datasets from FA patient samples demonstrated significant concordance with the *in vitro* dependency profiles. Second, intrinsic limitations of cancer-derived datasets e.g. mutational burden, chromosomal instability, and lineage-specific tumor evolution may introduce biases that are not representative of FA physiology. Third, while our study proposes biologically coherent mechanisms linking aldehyde metabolism, compensatory stress programs, and FA-like vulnerability states, definitive validation requires a systematic experimental program. Prospective validation in primary FA patient-derived hematopoietic cells, longitudinal transcriptomic profiling, and integration with clinical phenotypes will be critical.

In summary, this integrative approach allowed us to derive a prioritized list of potential rescue-associated genetic factors, laying the groundwork for downstream drug repurposing or targeted therapeutic prediction. The translational value of our findings ultimately depends on targeted experimental validation in physiologically relevant FA systems.

## ACKNOWLEDGMENT

The authors acknowledge the financial support from the King Abdullah International Medical Research Center (KAIMRC) Saudi Arabia (Grant: NRC25/118/25). The work was also funded by the Research Council of Finland (No. 351507 to ZT).

## Declaration of AI and AI-assisted technologies in the writing process

During the preparation of this work the authors used chatGPT to improve readability and language. After using this tool, the authors reviewed and edited the content as needed and took full responsibility for the content of the publication.

## Author Contributions

K.S. conceived and designed the study. K.S and Z.T performed the analyses, interpreted the data, and drafted the manuscript. All authors contributed to the interpretation of the results and critically reviewed the manuscript.

## SUPPLEMENTARY MATERIAL AND METHODS

## Supplementary Section S1

### Predictive modeling and feature ranking

Two complementary classification strategies were implemented: 1) Binary classification using low and high dependency groups (**Supplementary Table 1b, Figure 3)**, 2) multiclass classification using all three dependency groups (Low, Medium, and High) ((**Supplementary Table 1a**, **Supplementary Figure 3**). For both strategies, the dataset was randomly partitioned into 70% (training set) and 30% (independent test) using a stratified train-test split (random_state = 42). Stratified sampling preserved the class distribution across both subsets. The training set was used exclusively for model development and hyperparameter optimization, whereas the independent test set was reserved for unbiased performance evaluation. Three supervised machine learning algorithms were subsequently trained and compared to identify molecular features associated with FA pathway dependency (**Supplementary Table 1**).

Model performance was evaluated exclusively on the independent test dataset using multiple complementary metrics. Precision, recall, F1-score, and support were calculated for each dependency class using the classification report to assess class-specific predictive performance (**Supplementary Figure 3**). Confusion matrices were generated to visualize correctly and incorrectly classified samples across the low, medium, and high dependency groups. Receiver operating characteristic (ROC) curves and the corresponding area under the curve (AUC) values were computed from predicted class probabilities. For binary classification (scenario 1), ROC-AUC quantified discrimination between the low and high dependency groups. For multiclass classification (scenario 2), ROC curves were generated using a one-versus-rest strategy following label binarization, and both macro-averaged and weighted-averaged AUC values were calculated to evaluate overall classification performance across all dependency classes.

### S1.1: Random Forest

A Random Forest classifier comprising 200 decision trees (n_estimators = 200) was trained using the training dataset. Random Forest was selected because it effectively captures complex nonlinear relationships and feature interactions while remaining robust to high-dimensional, heterogeneous biological data and overfitting. During training, each tree was constructed from a bootstrap sample, with a random subset of predictor variables evaluated at each split to improve model diversity and generalization. The trained model was subsequently used to predict CRISPR dependency classes in the independent test dataset. Feature importance was quantified using the mean decrease in node impurity (Gini importance), enabling the identification and ranking of molecular features that contributed most strongly to classification performance.

### S1.2: XGBoost

An Extreme Gradient Boosting (XGBoost) classifier was independently trained using the same training dataset. XGBoost was selected for its ability to model complex nonlinear relationships while achieving high predictive performance through regularized gradient boosting. The algorithm sequentially constructs decision trees, with each tree correcting errors made by preceding trees to iteratively improve classification accuracy. Model training employed the multi-class logarithmic loss (mlogloss) objective with a fixed random seed (random_state = 42) to ensure reproducibility. The trained model was subsequently used to predict dependency classes in the independent test dataset. Feature importance scores were extracted from the trained XGBoost model to rank molecular variables according to their contribution to classification performance.

### S1.3: Soft Voting Ensemble

To determine whether integrating complementary machine learning algorithms improved predictive performance, a soft-voting ensemble classifier was constructed by combining the Random Forest and XGBoost models. Soft voting was selected because it aggregates class probabilities from multiple classifiers, leveraging their complementary strengths while reducing model-specific prediction errors and improving robustness. The ensemble averaged the predicted class probabilities from both models before assigning the final dependency class. The ensemble was trained using the same training dataset and evaluated exclusively on the independent test dataset. Model performance was assessed solely on the independent test dataset to provide an unbiased estimate of predictive accuracy.

### S1.4: SHapley Additive exPlanations (SHAP)

To improve model interpretability and identify biologically relevant predictors, SHapley Additive exPlanations (SHAP) were applied to the trained Random forest and XGBoost models. SHAP, a game theory-based approach, quantifies the contribution of each feature to individual model predictions. Global feature importance was summarized using the mean absolute SHAP value across all samples, with higher values indicating greater influence on model predictions. SHAP-based rankings complemented the feature importance estimates obtained from the RFRandom Forest and XGBoost models, providing a more interpretable assessment of key molecular predictors.

## Supplementary Section S2

### Gene expression analysis of Fanconi anemia patient samples

### *S2.1:* Bulk RNA-seq data analysis

Raw RNA-seq data were downloaded from the NCBI Sequence Read Archive (SRA) using the SRA Toolkit (prefetch and fasterq-dump). Paired-end FASTQ files were generated for each sample and subjected to quality assessment using FastQC, followed by summary reporting with MultiQC. Sequence reads were aligned to the human reference genome GRCh38 primary assembly (GENCODE Release 50) using STAR version 2.7.11b. The genome index was generated from the GRCh38 primary assembly FASTA file together with the matching GENCODE v50 primary assembly gene annotation (GTF) using a splice junction database overhang of 100, corresponding to the 101 bp read length. Alignments were produced as coordinate-sorted BAM files and indexed with Samtools version 1.21. Gene-level read counts were then generated from the aligned BAM files using HTSeq version 2.0.3 with exon features summarized by gene identifiers from the GENCODE v50 annotation. Individual count files were subsequently merged into a single raw count matrix containing all 9 samples, which served as the input for downstream differential gene expression analysis. All computational analyses were performed on the CSC Puhti high-performance computing environment using SLURM batch jobs. The raw read counts were normalized using TMM normalization method to generate the final normalized ReadCount.csv file.

### S2.2: Single-cell RNA-sequencing and pseudobulk differential expression analysis

Cell Ranger-generated filtered_feature_bc_matrix.h5 files from six Fanconi anemia patients and five healthy donors were imported into R using Seurat. Each file was treated as one biological sample, cell barcodes were prefixed with the sample identifier, and donor and condition information were added to the metadata.

Quality control was performed separately for each sample. Cells were retained when they had 200–7,500 detected genes, at least 500 UMIs, and no more than 20% mitochondrial transcripts. Doublets were identified within each sample using scDblFinder and removed. Cell numbers before filtering, after quality control, and after doublet removal were recorded.

Quality-controlled cells were merged and normalized using Seurat’s LogNormalize method with a scale factor of 10,000. The 3,000 most variable genes were used for scaling and PCA. Neighbour detection and clustering were performed using the first 20 PCs at a resolution of 0.4, followed by UMAP visualization by cluster, condition, and donor.

Cluster markers were identified using FindAllMarkers with the Wilcoxon rank-sum test. Positively enriched genes expressed in at least 10% of cells and with an average log2 fold change of at least 0.25 were retained. HSPC clusters were identified automatically and scored using canonical HSPC markers, including CD34, PROM1, SPINK2, GATA2, MEIS1, MECOM, HLF, HOXA9, LMO2, KIT, and FLT3. Scores were reduced when clusters showed erythroid, myeloid, lymphoid, or megakaryocytic marker expression.

The selection procedure combined marker expression, marker detection percentage, significant cluster-marker results, and donor representation. Additional marker-supported clusters were included when needed to ensure that every donor contributed at least 20 HSPC cells. The same selected cluster set was applied to all donors. Selected HSPCs were also compared with all other cells using FindMarkers, and marker tables, dot plots, UMAPs, cluster scores, and donor-coverage summaries were exported.

Raw UMI counts from selected HSPCs were summed by the donor to generate a pseudobulk matrix with one column per biological sample. Genes were retained when they had at least 10 total counts and were detected in at least three donors. Donor-level quality was assessed using library sizes, detected gene numbers, principal component analysis, and sample-correlation analysis.

Genome-wide differential expression was performed using DESeq2 with the design condition, using healthy donors as the reference. Positive log2 fold-change values indicated higher expression in FA. P-values were adjusted using the Benjamini-Hochberg method, and genes with adjusted p-values below 0.05 and absolute log2 fold changes of at least 1 were classified as differentially expressed. Log2 fold-change shrinkage using apeglm was applied when available.

Raw pseudobulk counts, DESeq2-normalized counts, log2-normalized counts, counts per million, variance-stabilized expression, genome-wide and target-gene differential-expression tables, volcano plots, quality-control results and R session information were exported.

### *S2.3:* Cross-platform concordance analysis

To evaluate the consistency of transcriptional responses across independent transcriptomic datasets, differential gene expression results from GEO microarray, bulk RNA-sequencing, and single-cell RNA-sequencing were compared independently with the DepMap CRISPR-derived gene prioritization dataset. Genes common to each pairwise comparison were matched by gene symbols. For each shared gene, the direction of regulation was determined from the sign of the log fold change (positive or negative). Genes showing the same direction in both datasets were classified as concordant, whereas genes with opposite directions were classified as discordant.

Pairwise agreement was quantified as the percentage of concordant genes among all common genes. Pearson’s correlation coefficient (r) was calculated to assess linear agreement between continuous log fold-change values, whereas Spearman’s rank correlation coefficient (ρ) and a direction-weighted Spearman correlation were used to evaluate monotonic relationships while accounting for gene ranking and regulation direction. The statistical significance of directional agreement was further assessed using a one-sided exact binomial test under the null hypothesis that concordant and discordant genes occur with equal probability (P = 0.5). A concordance network was generated using python program (NetworkX), where nodes represent individual datasets and edge labels summarize concordance percentage, number of shared genes, and correlation coefficients.

## Supplementary Section S3

### Computational workflow for LINCS-based transcriptional response analysis

### S3.1 LINCS perturbation dataset

Identification of compounds capable of modulating disease-associated molecular pathways requires large-scale transcriptomic datasets describing cellular responses to chemical perturbations. For this purpose, we utilized the Library of Integrated Network-based Cellular Signatures (LINCS) L1000 Level 5 dataset (GSE92742), one of the largest publicly available resources for systematically characterizing transcriptional responses induced by small-molecule compounds across a diverse collection of human cell lines ^17^. The LINCS L1000 platform directly measures the expression of 978 landmark genes and computationally infers the remaining transcriptome, enabling high-throughput characterization of compound-induced molecular responses while maintaining a manageable experimental scale.

Among the multiple processing levels provided by the LINCS consortium, the Level 5 dataset was selected because it contains consensus perturbation signatures generated using the Moderated Z-score (MODZ) algorithm, which integrates multiple biological replicates into a single robust transcriptional profile. Compared with individual replicate measurements (Levels 1-4), Level 5 signatures substantially reduce experimental variability while improving reproducibility, making them particularly suitable for large-scale transcriptomic analyses and compound prioritization. Together with the Level 5 expression matrix, the corresponding metadata describing perturbation signatures, compound identities, treatment conditions, perturbation types, cell lines, and landmark gene annotations were downloaded and incorporated into the computational workflow. Only chemical perturbation experiments (pert_type = trt_cp) were retained for downstream analyses, whereas genetic perturbations, ligand treatments, and other perturbation types were excluded to ensure that all analyzed signatures represented transcriptional responses induced by small-molecule compounds.

Rather than restricting the analysis to a single tissue type, the present study investigated transcriptional responses across two predefined groups of DepMap cell lines identified in our previous analyses. These groups represented distinct cellular phenotypes referred to as the low- and high-dependency groups. The low-dependency group consisted of 232 cell lines, whereas the high-dependency group comprised 84 cell lines (**Table S2**). Perturbation signatures generated within each group were extracted independently, enabling systematic comparison of compound-induced transcriptional responses between the two predefined cellular phenotypes while maintaining an identical marker gene panel and analytical framework.

Although the LINCS dataset provides transcriptome-wide perturbation profiles for hundreds of thousands of experiments, only a subset of genes was relevant to the biological question addressed in the present study (n=96, **S1**). Consequently, the next step involved constructing a Fanconi anemia (FA)-associated marker gene panel that could be consistently quantified across all selected perturbation signatures.

### S3.2 Construction of FA gene signatures

To characterize compound-induced transcriptional responses associated with Fanconi anemia, we assembled a panel of 96 FA-associated marker genes from our previous analyses (**S1**). The marker panel was designed to capture multiple biological processes implicated in FA and therefore included genes involved in aldehyde metabolism, oxidative stress regulation, lipid metabolism, cellular differentiation, immune regulation, signal transduction, cytoskeletal organization, cell adhesion, and additional pathways associated with disease biology. Members of the aldehyde dehydrogenase (ALDH) family constituted an important component of this marker panel because impaired aldehyde detoxification represents one of the major biological mechanisms contributing to DNA damage and disease progression in FA.

Because the LINCS L1000 platform directly measures only 978 landmark genes, all selected marker genes were matched with LINCS landmark gene set before downstream analyses. Mapping of the FA-associated marker panel demonstrated that 73 of the 96 marker genes (76.0%) were represented in the LINCS L1000 landmark gene set, whereas 23 genes (24.0%) were absent because they were not included among the measured landmark genes (**Supplementary Figure 7C**). This strategy minimizes uncertainty introduced by gene-expression imputation while maintaining direct comparability across all perturbation signatures included in the study.

The resulting marker panel represents a biologically informed transcriptional fingerprint of FA and serves as the common analytical framework throughout the study. Rather than focusing on individual genes independently, subsequent analyses evaluate coordinated transcriptional responses across the complete marker panel, thereby providing a systems-level representation of compound-induced perturbations. Normalized expression values corresponding to these genes were extracted from every selected LINCS perturbation signature to construct expression matrices suitable for downstream comparative analyses.

### S3.3 Generation of marker expression matrices

For every selected chemical perturbation signature, normalized MODZ expression values corresponding to the mapped FA marker genes were extracted from the LINCS Level 5 expression matrix. Each perturbation signature therefore produced a vector of normalized transcriptional responses describing the expression of the complete marker panel following compound treatment. The extracted expression profiles were subsequently organized into four complementary expression matrices representing different aspects of the transcriptional landscape. First, a collapsed marker expression matrix was generated by combining all selected perturbation signatures into a single dataset. This matrix provides a comprehensive overview of compound-induced transcriptional responses across the entire study population and serves as the basis for analyses investigating global transcriptional variation. Second, perturbation signatures belonging to the predefined low-dependency cell line group were combined to generate a low-dependency marker expression matrix. This matrix summarizes transcriptional responses observed exclusively within the low-dependency cellular phenotype. Third, perturbation signatures generated in the high-dependency cell line group were assembled into a corresponding high-dependency marker expression matrix, enabling independent characterization of compound-induced transcriptional responses within the second cellular phenotype. Finally, a differential marker expression matrix was constructed by calculating the difference between the average marker expression profiles of the low- and high-dependency groups (Low − High). This matrix directly highlights marker genes exhibiting the largest transcriptional differences between the two predefined cellular phenotypes and facilitates identification of compounds producing distinct responses across the two groups.

Together, these complementary matrices capture both the global and phenotype-specific transcriptional characteristics of the LINCS perturbation dataset. The collapsed matrix provides an overview of all perturbation signatures, whereas the low- and high-dependency matrices characterize transcriptional responses within each cellular phenotype. The differential matrix further emphasizes transcriptional changes that distinguish the two groups and therefore complements the individual group-specific analyses. Having established these expression matrices, exploratory transcriptomic analyses were next performed to characterize global expression patterns, investigate relationships among perturbation signatures, identify compounds exhibiting the strongest transcriptional responses, and evaluate similarities among highly responsive compounds. These analyses are described in the following sections.

### S3.4 Characterization of marker gene expression profiles

Heatmaps were generated to provide an overview of compound-induced transcriptional responses across the selected FA marker genes and to facilitate comparison between the low- and high-dependency cellular phenotypes **(Table S-S2)**. Because heatmaps enable simultaneous visualization of multiple genes and compounds, they provide an intuitive representation of transcriptional heterogeneity within large expression datasets. Visualizing every perturbation signature included in the LINCS dataset would substantially reduce interpretability due to the large number of compounds analyzed. Therefore, compounds were first ranked according to the variance of their marker gene expression profiles across the gene panel. The thirty compounds exhibiting the highest transcriptional variability were subsequently selected for visualization, as these compounds capture the greatest diversity of transcriptional responses within each dataset while maintaining a clear and interpretable figure.

For each expression matrix, marker gene expression values were displayed as normalized MODZ scores, where positive values indicate relative transcriptional upregulation and negative values indwebicate relative transcriptional downregulation compared with the LINCS reference distribution. Two independent heatmaps (**Figure 7A-B**) were generated representing the low-dependency and high-dependency group. This design enables direct visual comparison of compound-induced transcriptional patterns between the two predefined cellular phenotypes while simultaneously highlighting genes exhibiting the largest differences in expression. Although heatmaps effectively illustrate local expression patterns, they do not provide a quantitative summary of the overall magnitude of compound-induced transcriptional perturbations across the two predefined cellular phenotypes. Therefore, a perturbational activity score was calculated for each compound to enable quantitative comparison of transcriptional responses between the low- and high-dependency groups.

### S3.5 Quantification of perturbational activity and identification of transcriptionally responsive compounds

To compare the overall magnitude of compound-induced transcriptional perturbations between the low- and high-dependency cellular phenotypes, a perturbational activity score was first calculated for every perturbation signature. The perturbational activity score was defined as the mean absolute MODZ-normalized expression value across all mapped FA-associated marker genes. For compounds represented by multiple LINCS perturbation signatures, only the signature exhibiting the highest perturbational activity score was retained, thereby avoiding redundant representation of the same compound while preserving its strongest observed transcriptional response. Perturbational activity scores from the low- and high-dependency groups were subsequently compared using scatter plot visualization (**Figure 7C**). To identify compounds producing the strongest overall perturbation across the FA-associated marker panel, a transcriptional response score was calculated for every compound. Rather than evaluating individual marker genes independently, this approach summarizes the combined transcriptional response across the complete marker panel into a single quantitative measure, enabling systematic ranking of compounds according to the magnitude of their induced expression changes. Prior to score calculation, gene-wise MODZ expression values were standardized across compounds using z-score normalization. The transcriptional response score was then calculated as the Euclidean norm of the standardized marker gene expression vector:

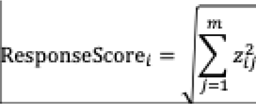

where *z_ij_* denotes the standardized expression value of marker gene *i*for compound *i*, and *m* represents the total number of marker genes included in the analysis. The Euclidean norm provides a robust measure of overall transcriptional perturbation because it simultaneously incorporates expression changes across all marker genes while remaining independent of the direction of regulation. Consequently, compounds exhibiting extensive transcriptional changes receive higher response scores regardless of whether individual genes are predominantly upregulated or downregulated, whereas compounds producing relatively small expression changes receive lower scores.

Following calculation of perturbational activity scores (**Figure 7C**), compounds were independently ranked according to their transcriptional response scores, and the twenty highest-scoring compounds were selected for further investigation (**Figure 7D**). These compounds represent the strongest global perturbagens within the selected FA-associated marker panel and were used for downstream analyses examining similarities and differences in their transcriptional response profiles. Although the response score identifies compounds producing the largest overall transcriptional perturbations, it does not indicate whether highly responsive compounds exhibit similar or distinct expression patterns. To investigate these relationships, correlation analysis was subsequently performed using the standardized marker gene expression profiles of the highest-ranked compounds.

### S3.6 Correlation analysis of highly responsive compounds

Whereas the perturbational activity score summarizes the average magnitude of transcriptional perturbation and the transcriptional response score quantifies the overall multivariate perturbation strength, neither metric captures similarities in gene-specific transcriptional response patterns among compounds. Therefore, correlation analysis was performed to investigate whether the highest-ranked compounds induce comparable or distinct transcriptional responses across the FA-associated marker panel.

The twenty compounds exhibiting the highest transcriptional response scores were selected for this analysis. Pairwise Pearson correlation coefficients were calculated using the standardized expression profiles of the 96 marker genes for each compound. Pearson correlation was chosen because it quantifies the degree of linear similarity between multivariate gene expression profiles while remaining independent of the overall magnitude of expression changes.

The resulting correlation coefficients were assembled into a symmetric correlation matrix and visualized as a clustered heatmap. Positive correlation coefficients indicate that two compounds produce similar patterns of marker gene regulation, whereas negative coefficients indicate opposing transcriptional responses. Compounds exhibiting strong positive correlations may therefore influence similar biological pathways or induce comparable transcriptional programs, whereas weakly correlated compounds may represent mechanistically distinct perturbations despite producing similarly large overall transcriptional responses.

This analysis complements the transcriptional response score by distinguishing compounds that generate comparable molecular signatures from those exhibiting unique expression profiles, thereby providing additional insight into the diversity of compound-induced transcriptional perturbations within the LINCS dataset.

### S3.7 Quality assessment of marker expression data

To ensure the robustness of downstream analyses and to characterize the statistical properties of the processed expression matrices, several complementary quality assessment procedures were performed using the complete marker expression dataset.

First, the overall distribution of normalized marker expression values was examined across all perturbation signatures. Because LINCS Level 5 signatures are represented as normalized MODZ scores, the global expression distribution was expected to be centered near zero. Inspection of this distribution enables identification of potential systematic biases or abnormal expression patterns that could influence subsequent analyses.

Second, gene-wise variance was calculated across all perturbation signatures to identify marker genes contributing most strongly to transcriptional heterogeneity. Genes exhibiting higher variance are expected to provide greater discriminatory power when distinguishing compound-induced transcriptional responses, whereas genes with consistently low variance contribute relatively little to overall dataset variability.

Third, the proportion of selected FA-associated marker genes represented within the LINCS landmark gene set was summarized to evaluate marker coverage. Because the L1000 platform directly measures only landmark genes, assessment of marker coverage provides an important indicator of how comprehensively the selected biological signature is represented within the experimental dataset.

To further characterize dataset consistency, gene-wise mean expression values and sample-wise standard deviations were calculated across all perturbation signatures. These descriptive statistics provide complementary information regarding the overall behavior of the selected marker panel and facilitate identification of potential outlier samples or unusually variable expression profiles.

Finally, the relationship between gene-wise mean expression and variance was examined to evaluate whether transcriptional variability depended systematically on average expression levels. This analysis provides an additional assessment of dataset stability and helps identify highly variable genes that may contribute disproportionately to the observed transcriptional landscape.

Together, these complementary quality assessment procedures provide a comprehensive overview of the processed LINCS expression dataset and demonstrate that the generated expression matrices possess appropriate statistical characteristics for downstream exploratory transcriptomic analyses.

**Table S1:** Marker genes.

| Marker Gene names |
| --- |
| ALDH1A1, ALDH1A2, ALDH1A3, ALDH1B1, ALDH1L1, ALDH1L2, ALDH2, ALDH3A1, ALDH3A2, ALDH4A1, ALDH5A1, ALDH6A1, ALDH7A1, ALDH8A1, ALDH9A1, ALDH16A1, ALDH18A1, AKR1C3, ADH5, MGLL, GPX4, CARMIL1, PFKFB2, CYB561, SHTN1, STAC3, TRIM47, ALOX5AP, ABCC3, RHPN2, ME1, PLPP2, RAP1GAP, LSP1, AIFM2, AGAP2, LAPTM5, GPAT3, OSGIN1, CD37, PTPN3, SHB, LAT2, EMP2, RPP25, ACAP1, MEF2C, MYO1D, EPS8, PERP, C1ORF54, FAM78A, GSTO2, SH2D4A, NBL1, TSPAN15, SPIRE2, NPAS2, KLF5, AFF3, PRKCZ, EPB41L4B, CORO1A, ITGA4, RIPK4, DAGLA, NHLRC1, DMTN, CDC42EP2, ANKRD29, DOCK2, DOCK8, PKP2, INPP5D, MAP4K1, PATJ, MET, ARHGEF6, ZC2HC1C, PHLDA2, PLBD1, ITGA3, CXCR4, SELENBP1, C6ORF141, PLEKHA6, MEIS1, FA2H, EFNA1, GPRC5B, FAM83H, C1ORF226, TUSC1, C3ORF52, ANKEF1, C19ORF33 |

**Table S2:**
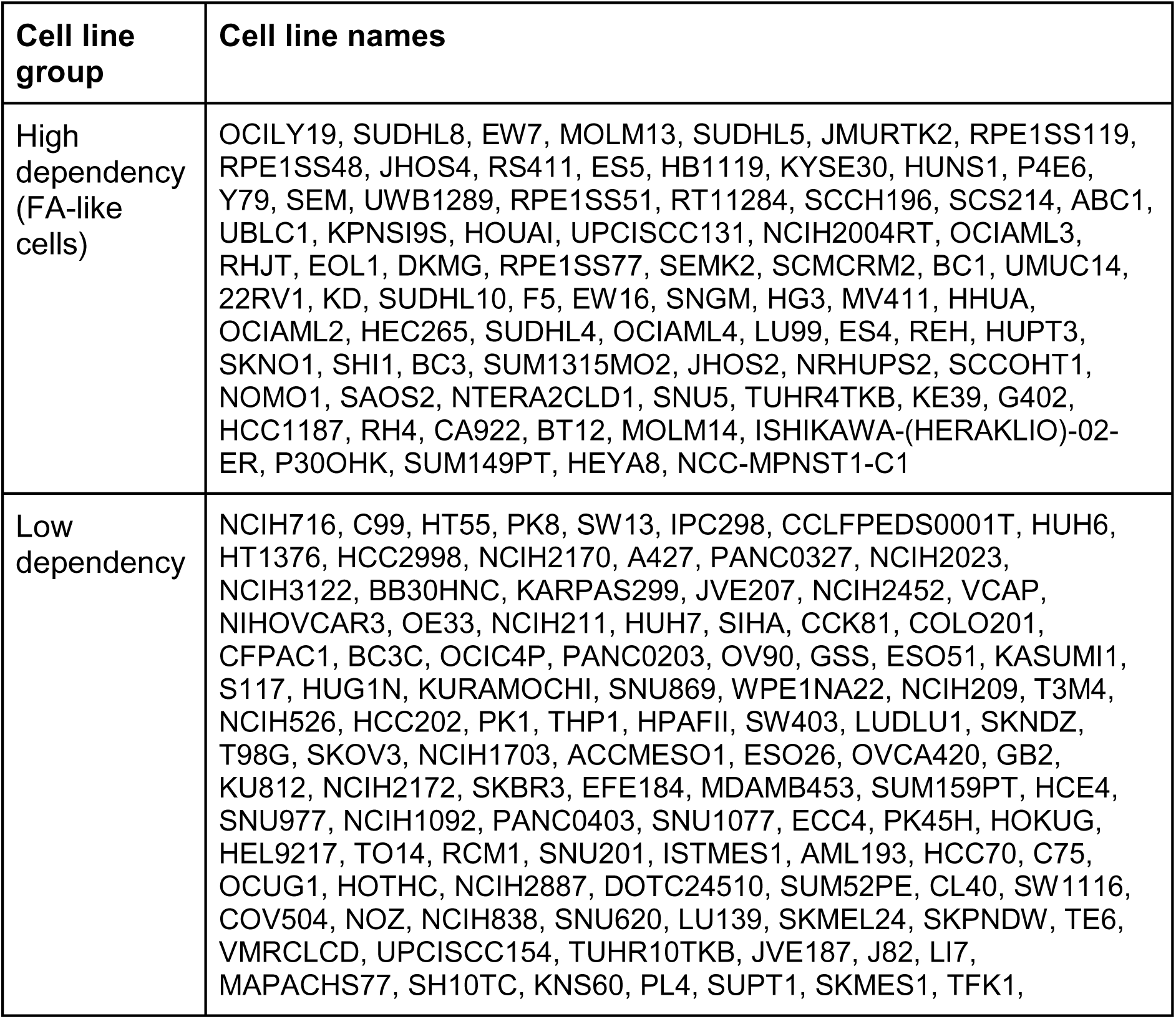
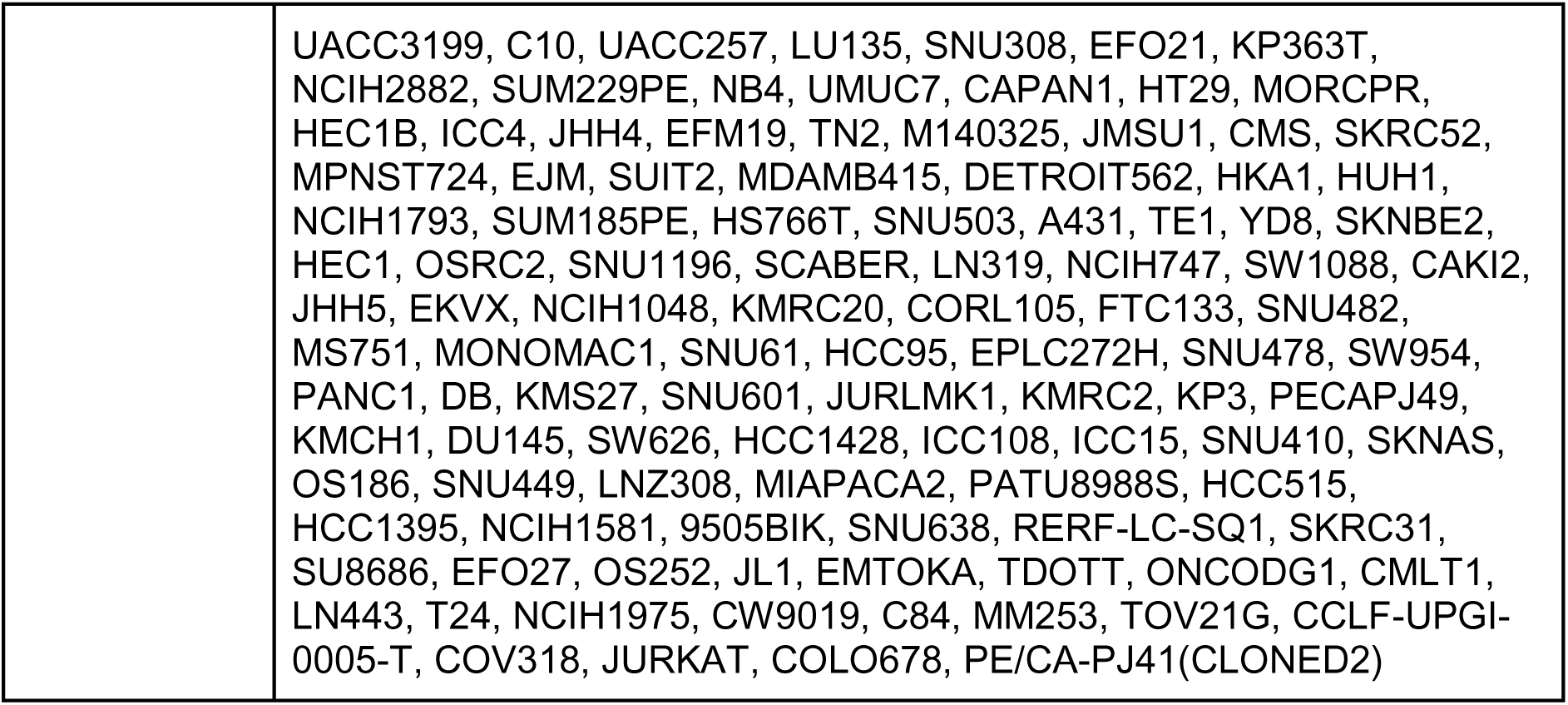
Two cell line groups related to FA.

## Additional Methods

### Pathway enrichment calculations

We utilize comprehensive, genome-wide datasets detailing protein interactions, localization, GO term annotation, co-expression (inverse), and pathway enrichment to evaluate the functional affiliation between specifying gene pairs. We wrote Python scripts that reads (1103 cell lines × ∼20k genes), filters to low vs high CRISPR dependency, computes ssGSEA pathway scores (per-sample pathway activities) using ‘gseapy’, tests pathway differences, and trains a simple classifier on pathway scores to find discriminative pathways (**Supplementary Figure 4B**). A separate run was performed using short listed candidate genes from t-test analysis with higher log fold change (LFC) mRNA (n=684). We performed statistical tests (t-test + FDR) to identify pathways that significantly differ (**Figure 4B**). We trained a sparse logistic regression on pathway activity scores to find a small multivariate signature (combination of pathways) that discriminates between the groups. In addition, curated gene lists were submitted to web-based gene-set enrichment analysis platform Enrichr (Ma’ayan Lab) against curated functional libraries including Gene Ontology (GO), Reactome to investigate their functional and biological relevance. Enrichment analyses were performed against multiple curated gene-set libraries, Significantly enriched biological processes and pathways were ranked using Enrichr’s combined score and adjusted *P* values to identify molecular functions and signaling networks associated with the prioritized genes ^43,44^.

### SHAP Interaction Analysis and Class-Specific Bias Testing

XGBoost was trained to distinguish low vs high dependency cell lines using RNA expression features. SHAP interaction values were computed using TreeExplainer to quantify pairwise feature interactions contributing to model predictions (**Supplementary Figure 5C**). For each gene pair, mean absolute SHAP interaction strength was calculated across all samples and stratified by class. Positive interaction values indicate synergistic gene effects, whereas negative values indicate antagonistic or compensatory relationships. Differences between low and high interaction distributions were evaluated using the Mann-Whitney U test, and interaction bias was quantified as the difference in mean interaction magnitude between classes. Gene pairs were ranked by class-specific interaction bias, and the most discriminatory interactions were visualized with annotated cell-line-level contributions.

### Top FA dependency ranking in Hematopoietic lineage

CRISPR dependency scores for FANCA, FANCC, and FANCG were extracted and summarized across cell lines using the mean FA dependency score (FA_mean) for **Figure 5D and Supplementary Figure 5D-E**. Cell lines were ranked from lowest to highest mean FA, with lower Chronos values indicating higher dependency. RNA expression differences were evaluated between the top FA-dependent cell lines and the remaining cell lines. For each RNA feature, group-level differences were assessed and visualized to identify genes showing lineage- or dependency-associated expression shifts.

### Dimensionality Reduction and Cluster Analysis of prioritized gene signature

To determine whether this prioritized gene signature captured biologically meaningful variation rather than statistical noise, we evaluated its ability to distinguish transcriptional relationships across a diverse panel of cancer cell lines. Expression values of the 61 concordant genes were extracted from all eligible cancer cell lines and standardized using Z-score normalization to ensure equal contribution of each gene to downstream analyses. Principal Component Analysis (PCA) was first performed to identify the major sources of global transcriptional variation captured by the selected gene signature. Because PCA is a linear dimensionality reduction approach, Uniform Manifold Approximation and Projection (UMAP) was additionally applied using 15 nearest neighbors and a minimum distance of 0.1 to preserve nonlinear relationships and local neighborhood structure among samples.

Cell lines were annotated according to both tissue lineage and FA dependency class (Low or High). Two-dimensional embeddings generated by PCA and UMAP were visualized to assess clustering patterns used in (**Figure 5A, Supplementary Figure 5A**). To quantitatively evaluate whether the observed group separation exceeded random expectation, pairwise euclidean distance matrices were calculated from the PCA coordinates, and Permutational Multivariate Analysis of Variance (PERMANOVA) with 999 permutations was performed independently for FA dependency class and tissue lineage. Statistical significance was determined using permutation-derived *P*-values. Effect sizes were reported as pseudo-R² values, reflecting the proportion of variance explained by each grouping factor.

**Supplementary Figure 1:**
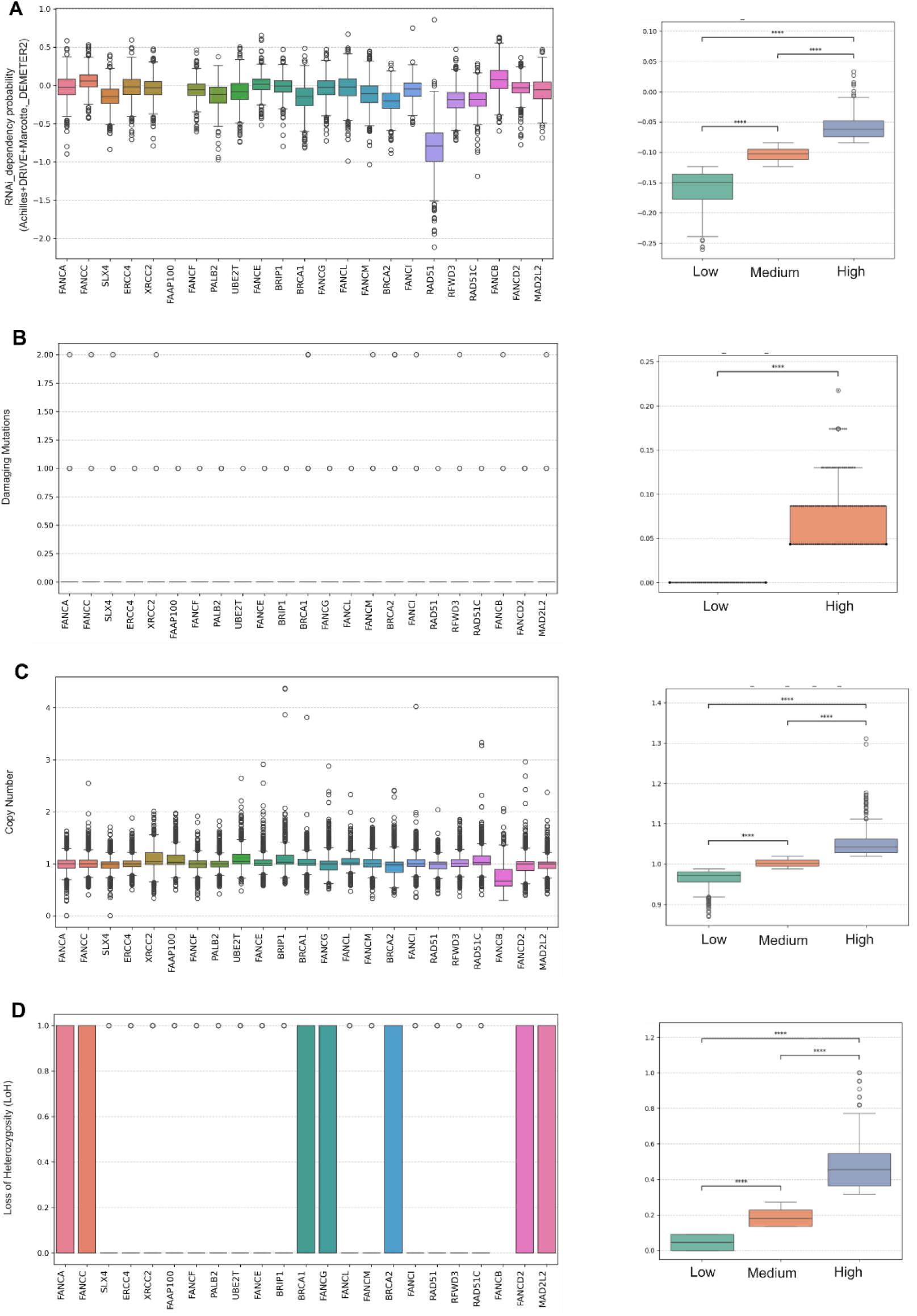
Molecular characteristics of combined FA genes. The boxplot illustrates the combined dependency/features of 23 FA genes across a panel of over 1000 cancer cell lines from diverse tissue origins, analyzed using the DepMap portal (https://depmap.org/portal). **A)** RNAi scores. **B)** Frequency of damaging mutations. **C)** Copy number variations (CNVs). **D)** loss of heterozygosity (LOH). Left panel: individual genes. Right panel: combined genes and classify based on low, medium, and high scores. Statistical significance was determined using t-test. Significance levels are denoted as P < 0.05 (*), P < 0.01 (**), and P < 0.001 (***); ns = not significant.

**Supplementary Figure 2:**
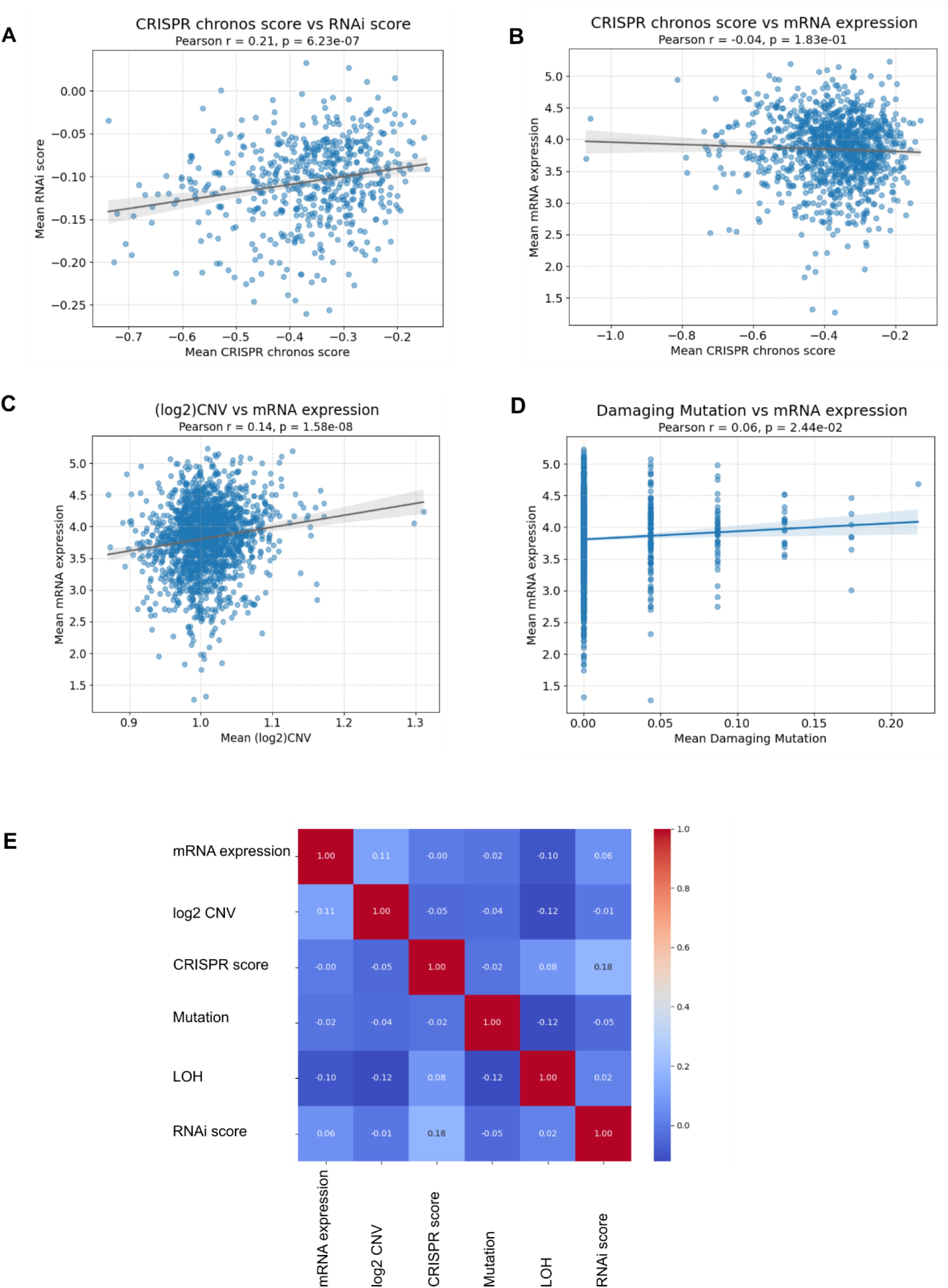
Pearson correlation of variable molecular signatures. Mean value 23 FA genes for each parameter using all available cell lines data was calculated and visualised through scatter plots **A)** CRISPR vs RNAi. **B)** CRISPR vs mRNA expression. **C)** CNV vs mRNA. **D)** mRNA vs damaging mutations. **E)** Heatmap exhibiting Pearson correlation of combined molecular signatures, annotated by color code. Red represents the highest correlation while dark blue highlights the lowest association.

**Supplementary Figure 3:**
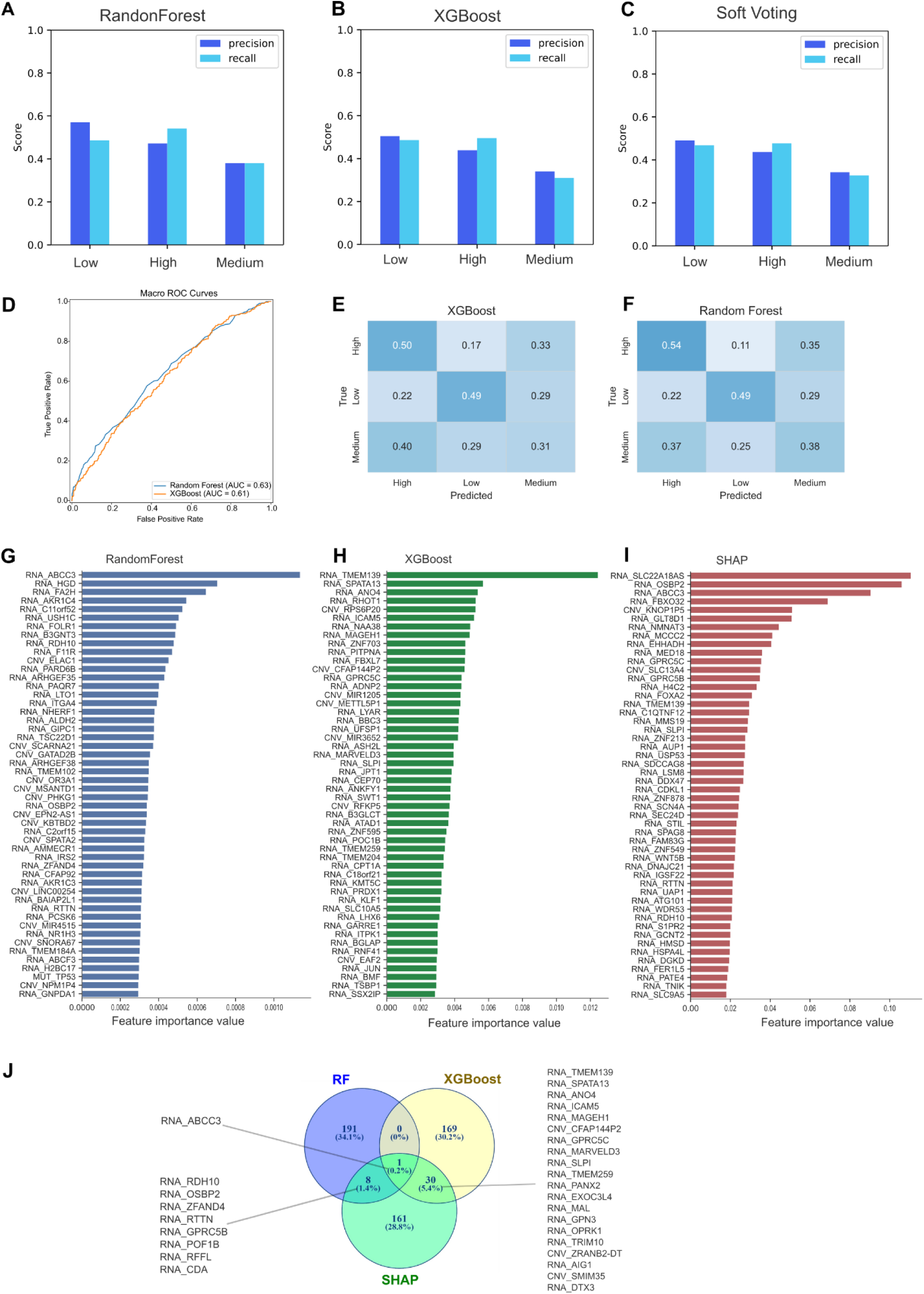
Multi-omics integration and FA dependency analysis in low, medium and high dependency group. **A)** Precision and recall for each dependency class obtained from these models using the independent testing dataset. Precision represents the proportion of predicted samples belonging to a class that were correctly classified, whereas recall represents the proportion of true samples successfully identified **A)** for Random Forest (RF), **B)** XGboost (XGB) classifier and **C)** soft voting ensemble. **D)** Receiver Operating Characteristic (ROC) curve was computed and plots the true positive rate (sensitivity) against the false positive rate (specificity) based on predicted class probabilities (low, high dependency groups), and performance was summarized using Area Under the Curve (AUC), with higher values indicating improved classification. **E)** Confusion matrices evaluated per-class accuracy summarizes the number of true positives, false positives, false negatives, and true negatives for each class by comparing predicted labels against true labels for RF, **F)** XGB. **G)** Molecular features were ranked according to their contribution to model prediction using the intrinsic feature importance scores calculated by each algorithm. The top-ranking variables are shown in descending order of importance. G) Random Forest importance reflects the average reduction in Gini impurity across all decision trees, whereas **H)** XGBoost importance represents the cumulative contribution of each feature during gradient-boosted tree construction. **I)** Mean absolute SHAP values were calculated from the trained XGBoost model using the independent testing dataset. Larger SHAP values indicate variables exerting greater influence on model predictions irrespective of direction. Features are ranked from highest to lowest overall contribution. **J)** Venn-diagram show overlapping genes identified in each predicting models.

**Supplementary Figure 4.**
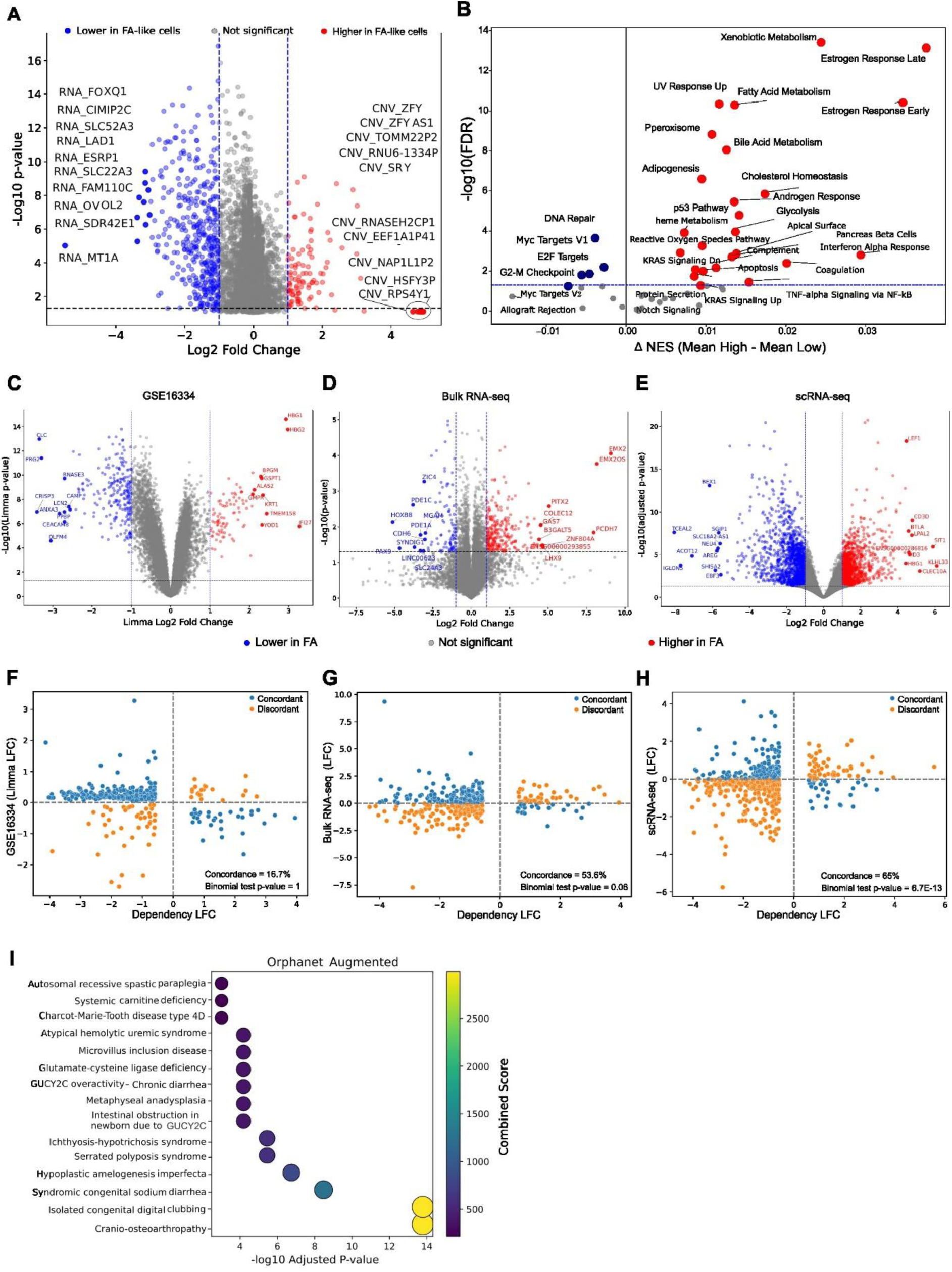
**A)** Volcano plot represents differential molecular features measured between low vs high CRISPR dependent groups. Each point represents an individual molecular feature (RNA expression, copy number variation (CNV), or mutation). A total of 17,123 molecular features were analyzed, including 5,364 mRNA expression, 11,758 CNV, and 1 mutation. Features with median values below 0.1 in both groups were excluded. Features located within the fold-change threshold are shown in gray, while the rest are shown in blue and red, while the top annotated features are highlighted and labelled. **B)** Pathway enrichment score (per-sample pathway activities) using GSEApy based on all the differential mRNA expression across low and high CRISPR dependencies. **C-E)** Volcano plot represents differentially expressed mRNA (Limma LFC) between **C)** normal volunteers and patients with Fanconi anemia. **D)** represents genome-wide differential expression performed against healthy donors and FA-patients using DESeq2. Positive log2 fold-change values indicated higher expression in FA. P-values were adjusted using the Benjamini-Hochberg method, and genes with adjusted p-values below 0.05 and absolute log2 fold changes of at least 1 were classified as differentially expressed. Log2 fold-change shrinkage using apeglm was applied when available. **E)** represents genome-wide differential expression performed against healthy donors and FA-patients from the bulk RNA-seq data. **F-H)** Scatter plots show the relationship between log fold-change (LFC) values for genes shared between cross three platform datasets vs LFC calculated from low versus high CRISPR dependency data, **F)** GEO microarray, **G)** bulk RNA-seq, and **H)** single-cell RNA-seq datasets. Each point represents one common gene. Concordance was evaluated based on the direction of regulation, where genes exhibiting the same sign of LFC were classified as concordant and those with opposite signs as discordant. Statistical significance of directional agreement was assessed using a one-sided exact binomial test. **I)** 41 concordant lower expressed genes were submitted to web-based gene-set enrichment analysis platform Enrichr (Ma’ayan Lab) against Orphanet augmented pathway.

**Supplementary Figure 5.**
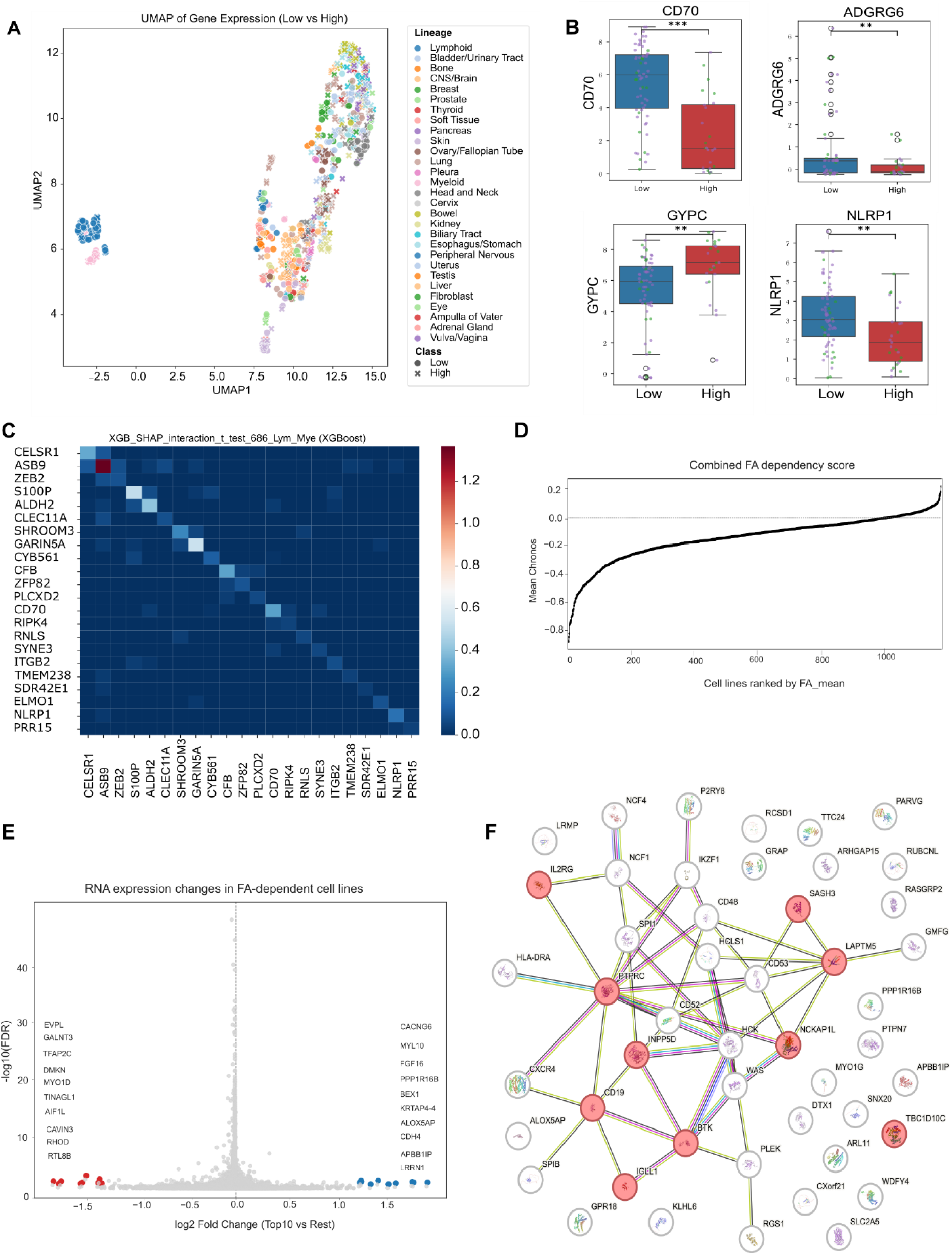
Hematopoietic lineage displays characteristic and predictable traits. **A)** UMAP showed coordinated lineage-specific transcriptional programs as distinct clusters. Samples are colored according to tissue lineage and shaped according to FA dependency class (Low or High). **B)** Boxplot highlighted top ranked differential expressed genes included CD70, GYP, ADGRG6 and NLRP1. colors are indicating respective lymphoid/myeloid lineages within the plot. Statistical significance was determined using t-test. Significance levels are denoted as P < 0.05 (*), P < 0.01 (**), and P < 0.001 (***); ns = not significant. **C)** SHAP interaction values were computed using TreeExplainer to quantify pairwise feature interactions contributing to model predictions. For each gene pair, mean absolute SHAP interaction strength was calculated across all samples and stratified by class. Positive interaction values (red) indicate synergistic gene effects, whereas negative values (blue) indicate antagonistic relationships. Gene pairs were ranked by class-specific interaction bias, and the most discriminatory interactions were visualized with annotated cell-line-level contributions. **D)** Cell lines were ranked by mean FA dependency scores of FANCA, FANCC, and FANCG. **E)** Volcano plot showed RNA expression differences between the top four FA-dependent cell lines versus remaining cell lines. **F)** Top 50 differential RNA of the four FA-dependent cell lines were subjected to enrichment networks using https://string-db.org/ with the high confidence threshold.

**Supplementary Figure 6.**
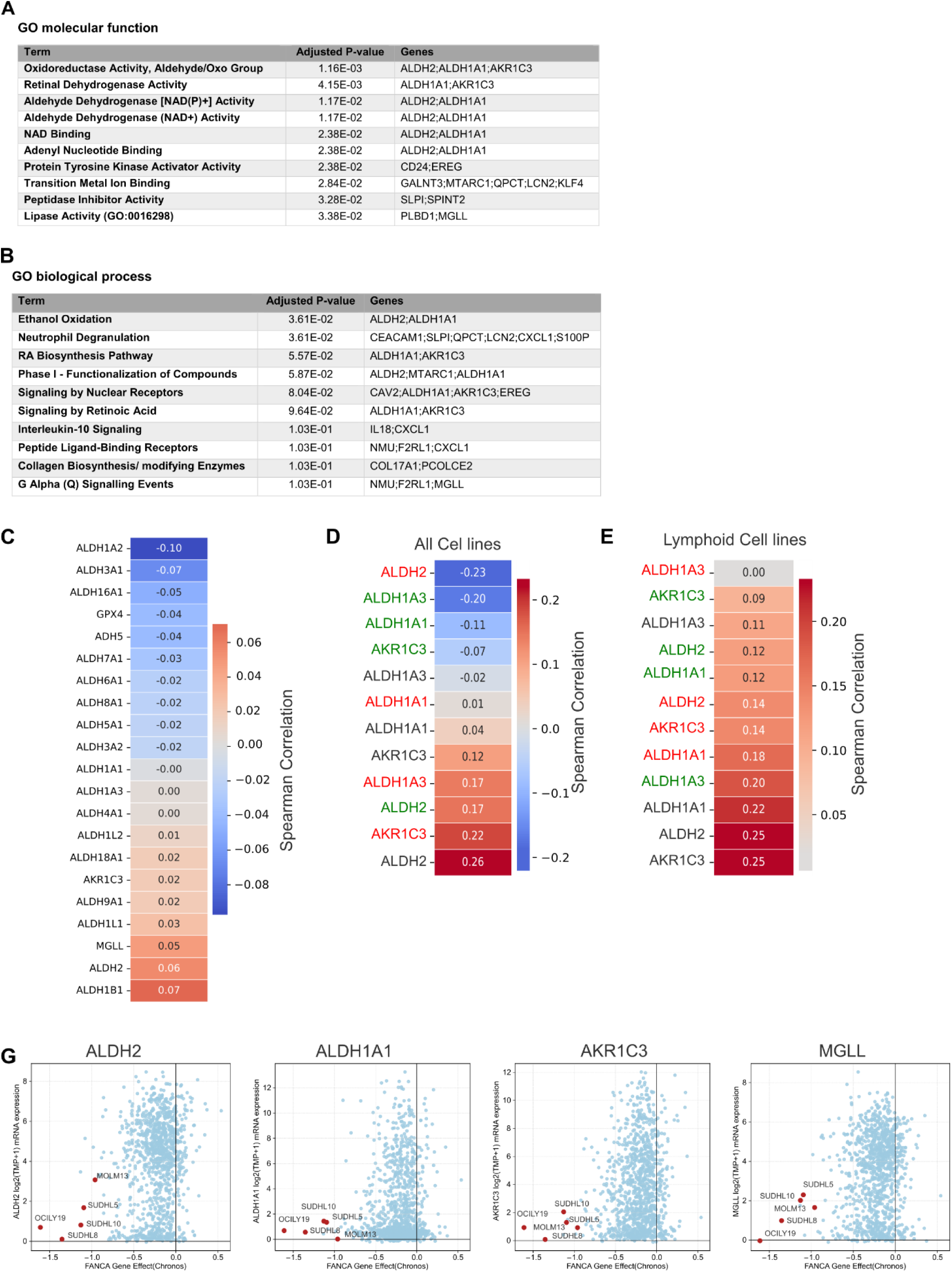
Endogenous aldehydes and Fanconi anaemia pathway interactions. **A)** Aldehyde metabolism and related pathway genes were identified redundant in pathway enrichment analysis, GO molecular function. **B)** GO biological process. **C)** Association of aldehyde dehydrogenase family members and extended partners were analysed for mRNA expression and CRISPR scores by Spearman correlation. **D)** Individual members and their expression were plotted against all stratified cell line’s median CRISPR score across all cell lines, **E)** in hematopoietic cell lines only. Font color code of the cell line: Black = all cell lines, Red = high dependency, Green= low dependency. **F)** Scatter plots illustrating the expression of four key genes involved in aldehyde metabolism, together with the FANCA CRISPR gene-effect score, identified a distinct subgroup of cell lines exhibiting low gene expression and low CRISPR dependency. Four lymphoid cell lines (OCILY-119, SUDHL-5, SUDHL-8, and SUDHL-10) and one myeloid cell line (MOLM-13), highlighted in red, consistently clustered within this low-expression, high-dependency group.

**Supplementary Figure 7:**
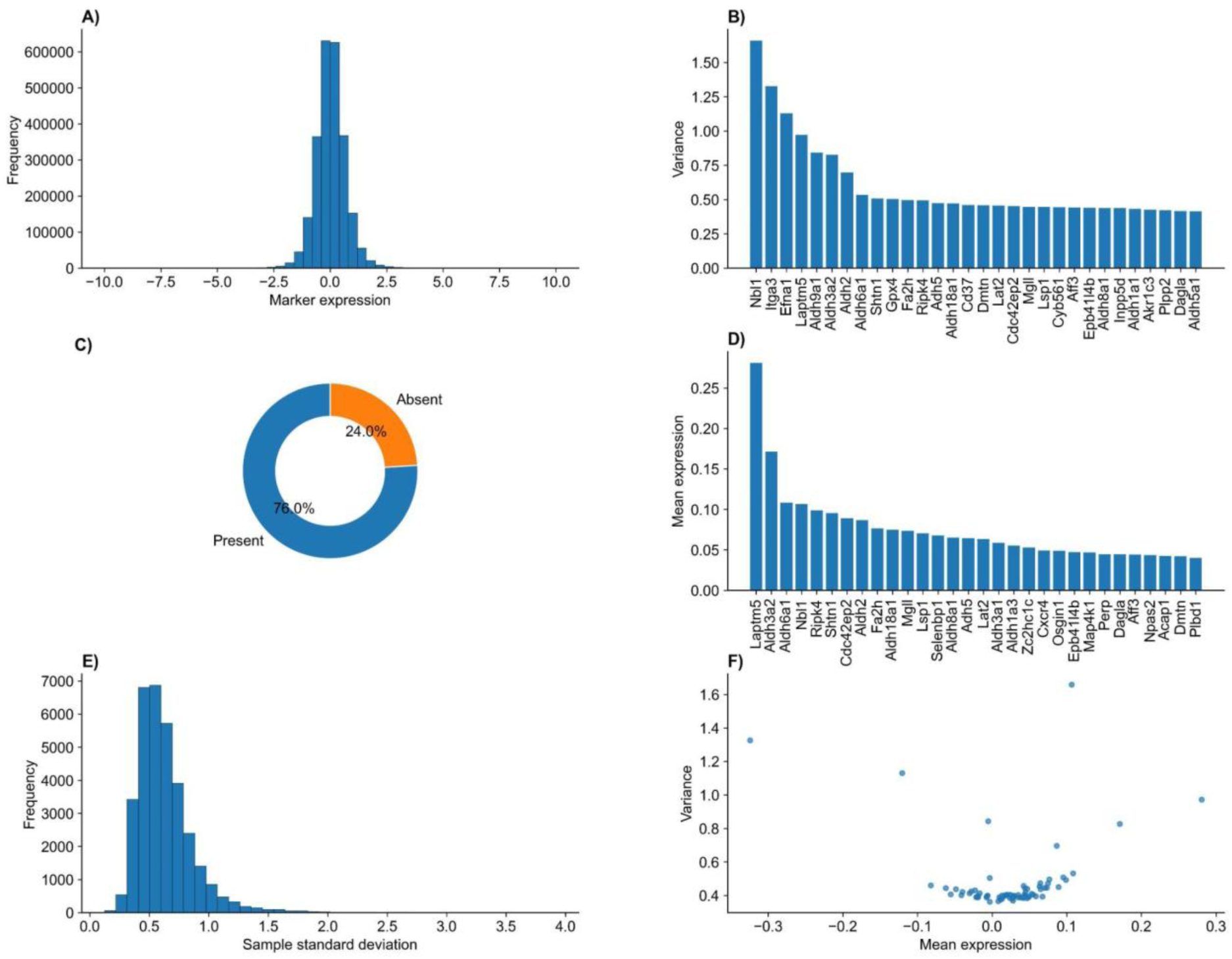
Quality assessment and descriptive statistics of LINCS marker expression data. Overview of the distribution and statistical characteristics of the marker gene expression dataset used in this study. **(A)** Distribution of standardized marker gene expression values across all compounds and marker genes. **(B)** Top 30 marker genes ranked by expression variance, highlighting genes contributing most to transcriptional heterogeneity. **(C)** Proportion of selected marker genes represented in the LINCS L1000 landmark gene set, showing the percentage of genes available for downstream analysis. **(D)** Top 30 marker genes ranked by mean expression across all compound profiles. **(E)** Distribution of sample-wise standard deviations, illustrating the variability of transcriptional responses among compounds. **(F)** Relationship between mean expression and variance for all marker genes, demonstrating the overall expression-variability characteristics of the dataset.

